# Reward revaluation biases hippocampal replay content away from the preferred outcome

**DOI:** 10.1101/397950

**Authors:** Alyssa A. Carey, Youki Tanaka, Matthijs A. A. van der Meer

**Author notes:** Correspondence should be addressed to MvdM, Department of Psychological and Brain Sciences, Dartmouth College, 3 Maynard St, Hanover, NH 03755.

## Abstract

The rodent hippocampus spontaneously generates bursts of neural activity (“replay”) which can depict spatial trajectories to reward locations, suggesting a role in model-based behavioral control. A largely separate literature emphasizes reward revaluation as the litmus test for such control, yet the content of hippocampal replay under revaluation conditions is unknown. We report that the content of awake hippocampal sharp wave-ripple events is biased away from the preferred outcome following reward revaluation, challenging the idea that such sequences reflect recent experience or trajectories toward the preferred goal.

## Main text

The spatial navigation literature has identified a role for the hippocampus in flexibly navigating toward goal locations. This behavior is thought to rely on knowledge of the environment (a “cognitive map”) to flexibly generate and decide between possible trajectories, an example of model-based behavioral control ^1^. Sequences of hippocampal place cell activity, colloquially referred to as “replay” when coincident with sharp wave-ripple complexes (SWRs) ^2, 3^, are a candidate neural underpinning for such planning, as suggested by the bias of such sequences toward goal locations ^4^ and behavioral impairments resulting from SWR disruption ^5^. In contrast, the conditioning literature operationalizes model-based control as sensitivity to reward revaluation ^6^. Reward revaluation causes rate remapping of hippocampal place cells ^7^ and hippocampal damage impairs some behaviors that rely on the discrimination of internal motivational states such as hunger and thirst ^8–10^. However, it is unknown how reward revaluation affects the content of hippocampal SWRs. A goal-directed bias in hippocampal SWRs predicts that its content favors the currently highly valued outcome, especially following a recent revaluation. Conversely, if hippocampal SWR content is unaffected by reward revaluation, this would call into question a role for SWR activity in model-based control.

To determine how hippocampal SWR content is affected by reward revaluation, we recorded neural ensembles from the dorsal CA1 area of the hippocampus as rats (n = 4, male) performed a T-maze task, which offered free choice between food (left arm) and water (right arm) outcomes (Figure 1a). The crucial manipulation in the experimental design was that prior to daily recording sessions, animals were alternately food- or water-restricted, revaluing the food and water outcomes due to a shift in their motivational state. As a result, animals exhibited a clear overall behavioral preference for the restricted outcome (Figure 1b; .90 ± .07 (SEM) food choices on food-restricted days, .19 ± .09 food choices on water-restriction days). To assess the statistical significance of this result, as well as all other major results in the study, we compared the observed difference between food- and water-restriction days to a resampled distribution of differences obtained by randomly permuting food- and water-restriction labels (*fr* and *wr*; gray bars in Figure 1b show the mean ± SD of this shuffled distribution). For this behavioral preference data, the observed difference (*diff*) was clearly larger than the shuffled differences (*z* = 4.32, *p* < .001); the positive z-score indicates that motivational state shifts behavioral preference *towards* the restricted outcome. Across trials within recording sessions, this preference was most clearly apparent from the second trial onward (Figure 1c). Although choice preference for the restricted outcome did not differ significantly between food and water-restricted days (.90 and .81 respectively, *p* = 0.12), trial length was shorter on food-restricted compared to water-restricted sessions (14.9 ± 5.2 s and 22.5 ± 7.5 s, *p* = .011). This indicates that on average, food-restricted animals were more motivated to work for food than water-restricted animals were in working for water, consistent with previously reported asymmetric effects of both restriction types ^11^.

**Figure 1:**
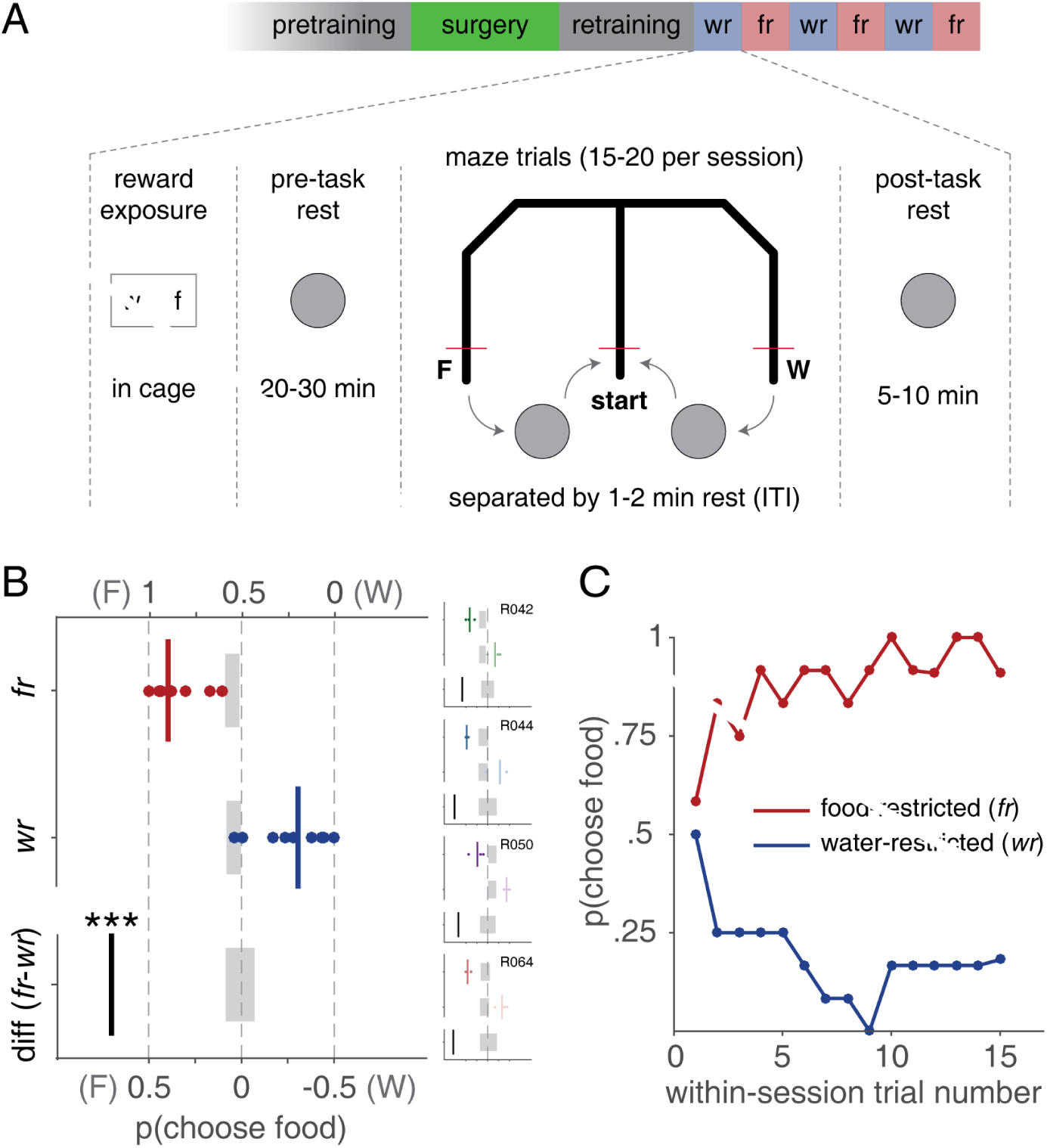
Experimental design, task and behavior. **A**: Rats (n = 4) experienced daily sessions on a T-maze offering a choice between food (F) at the end of the left arm and water (W) at the end of the right arm. Blocked trials ensured adequate sampling of each outcome (see *Online Methods* for details). Daily recording sessions consisted of exposure to both food and water outcomes in the home cage, followed by a pre-task rest session on a movable platform. 15-20 T-maze trials, separated by a 1-2 minute intertrial interval (ITI) were performed next, followed by post-task rest. Crucially, across recording sessions, rats were alternately water-restricted (blue, *wr*) or food-restricted (red, *fr*). **B**: Proportions of food (left) arm choices on free-choice trials (shown on the upper horizontal axis). Following food restriction (*fr*, red) rats preferred the food (F, left) arm, whereas following water restriction (*wr*, blue) the water (W, right) arm was preferred. As a result, the difference in choice proportions (*diff*, food-minus water-restriction, shown on the lower horizontal axis) was strongly biased towards the food (left) arm. Dots indicate single sessions, vertical bar the mean across sessions. Asterisks indicate significance level (*^∗^*: *p* < .05; *^∗∗^*: *p* < .01; *^∗∗∗^*: *p* < .001) by comparison with a resampled distribution based on shuffling food and water restriction labels (gray bar center indicates the mean, width indicates the standard deviation across shuffles). Horizontal axis direction is reversed (positive numbers on the left) so that the location of the food arm in the plots matches its physical location on the maze (both on the left). Overall, rats chose both arms a comparable number of times (193 food, 197 water). Insets show choice proportions for each individual subject. **C**: Within-session choice proportions of choosing the food arm following food restriction (red) and water restriction (blue), averaged across all sessions and subjects. Note that these proportions do not necessarily sum to one because food and water restriction sessions occurred on different days.

Our neural analyses focused on SWR-associated ensemble activity patterns (“replay”) ^2, 3^. Specifically, we sought to test how the *content* of such patterns – i.e. left (food) or right (water) trials on the T-maze – related to the animals’ motivational state and behavior. To access SWR content, we applied two different analyses using Bayesian decoding ^12^ (see Figure S1 for a schematic of the analysis pipelines). The first, *sequenceless decoding* analysis, started with the identifiction of putative SWR events based on spectral properties of the local field potential (LFP) and multi-unit firing activity (MUA). Next, we decoded each putative SWR event as a single time window, and computed its log odds of corresponding to a left or right trial relative to a shuffled distribution (Figure S2 and *Online Methods*; Figure S3 shows that left and right trial neural activity was clearly distinguishable during running on the track). Thus, this analysis does not involve sequential structure in neural activity. In addition to reporting unthresholded log odds across all SWR events, we also used a significance threshold to identify and retain only those events that clearly decoded to left or right trials. For the second analysis, *sequence detection*, we first decoded the full session data using a 25 ms moving window, and then applied sequence detection ^4^ to identify spatiotemporal sequences that decoded to either the left or right trajectory on the track (but not both, *Online Methods*). Only sequences that passed a comparison against a chance distribution based on shuffled neural activity, and that overlapped with a putative SWR event (defined as above), were retained for subsequent analysis.

### **Reward revaluation shifts sequenceless SWR content** away from the preferred outcome

To determine the relationship between motivational state and SWR content, we first quantified SWR content by decoding entire SWR events as a single time window, and z-scoring the log odds of the left vs. right posterior 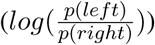 relative to a resampled distribution obtained by randomly permuting left and right arm tuning curves (sequenceless decoding; see Figure S2 and *Online Methods*). Plotting the resulting log odds z-score across recording sessions revealed a striking pattern: motivational shifts biased SWR content *away from* rats’ behavioral preferences (Figure 2a). That is, for food restriction sessions (*fr*, red shading in Figure 2a) rats preferentially chose the food (left) arm (black line), but SWR content was biased toward the water (right) arm (green line). For water restriction sessions (*wr*, blue shading), rather than shifting SWR content more towards the water (right) arm, SWR content shifted in the direction of the food (left) arm. Thus, the difference (*diff*) in SWR content log odds z-scores between food and water restriction was significantly negative, indicating a shift in the opposite direction from the restricted outcome (Figure 2b; food restriction –.42 ± .27, water restriction .40 ± .28, difference *z* = –2.68, *p* = .0073 compared to a distribution in which motivational states were randomly shuffled). Importantly, each individual subject exhibited this pattern (Figure 2b, insets). Moreover, a linear mixed model with subject-specific intercepts and SWR content log odds z-score as the dependent variable was clearly improved by the addition of motivational state (likelihood ratio test, *LRstat*_Δ*df* =1_ = 9.3, *p* = .0023). Similarly, behavioral choice was strongly anticorrelated with SWR content across sessions (Pearson’s *r*_(19)_ = –.75, *p* < .001; see Figure S4 for individual subject data).

**Figure 2:**
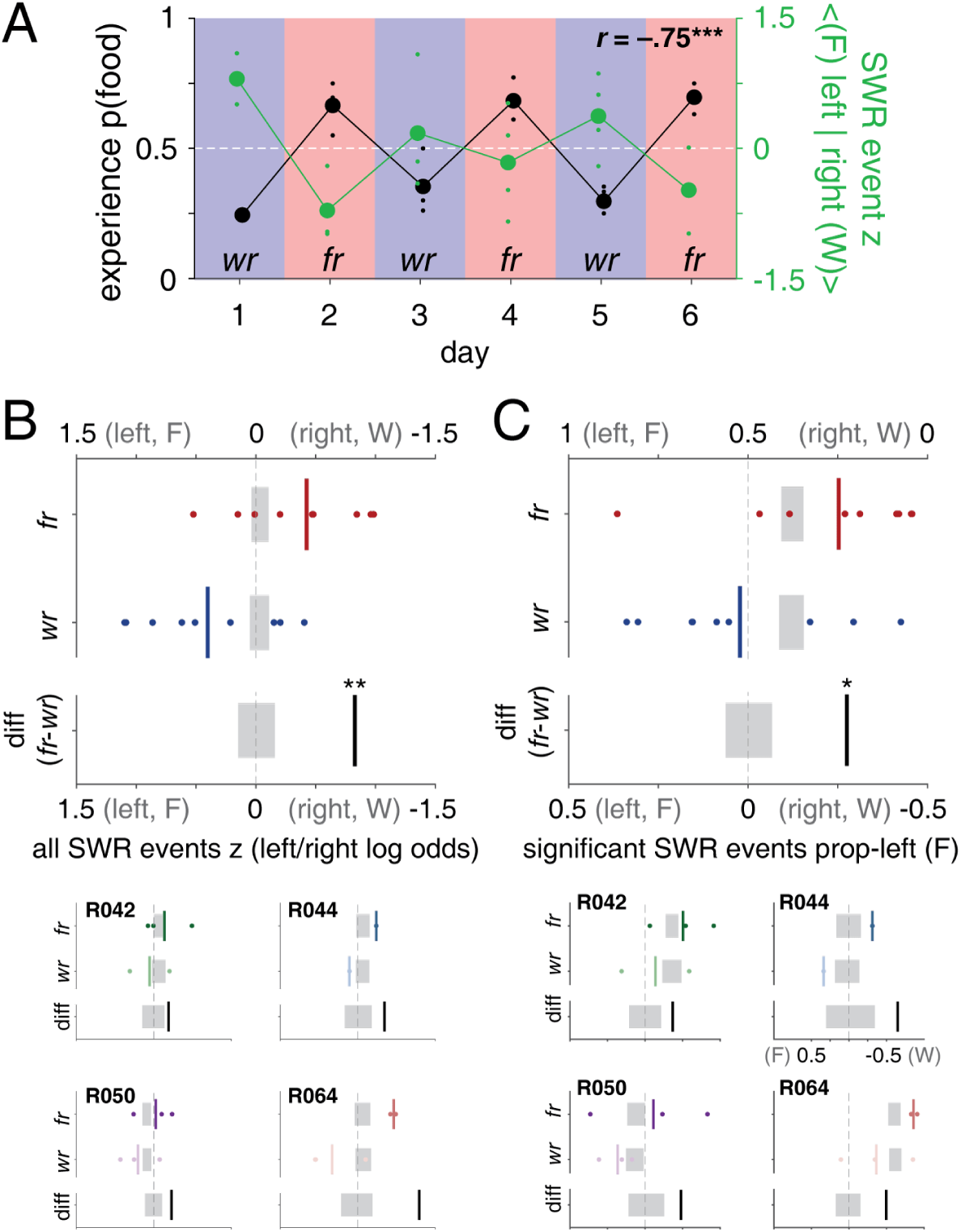
Sequenceless SWR content is biased in the opposite direction from motivational shifts. **A**: Motivational state changes the Z-scored log odds (left vs. right; recall that the left arm always leads to food and the right arm always leads to water) of decoded SWR content (green) opposite to behavioral preference (probability of choosing the food arm, black), as indicated by the negative correlation (*r* = –.75, *p* < .001). Positive z-scores correspond to left (food) trials, negative z-scores correspond to right (water) trials. Shaded areas indicate water- (blue, *wr* and food-restriction sessions (red, *fr*). Large markers show averages across subjects, small markers indicate single recording sessions. **B**: Mean z-scored log odds across food-restriction sessions (red), water-restriction sessions (blue; upper horizontal axis), and their difference (*diff*, black; lower horizontal axis). Dots indicate single sessions, vertical bar shows the mean across sessions. Asterisks indicate significance level (*^∗^*: *p* < 0.05, *^∗∗^*: *p* < 0.01, *^∗∗∗^*: *p* < 0.001) by comparison with a resampled distribution based on shuffling food and water restriction labels across sessions (gray bar width indicates standard error of the mean across shuffles). Note that the observed difference in SWR content is significantly different from the shuffled distribution, and in the opposite direction compared to behavior (compare Figure 1b). **C**: Proportion of significance-thresholded SWR events that decode to the left (food) arm for food-restriction sessions (red), water-restriction sessions (blue; upper horizontal axis), and the difference between them (food restriction minus water restriction session proportions; bottom horizontal axis). Small panels in B and C show single-subject data; axis labels are omitted when identical to those in the average plots.

The above analysis included decoding of *all* candidate SWR events, but not all observed SWR events decode clearly to any single environment or location ^13^. Thus, it is possible that the above effects are due to an effect of motivational state on those events that do not in fact clearly decode to one arm or another. To address this issue, we thresholded the z-scored log odds to retain only those events that clearly favored left or right (i.e. that exceeded *p* = .05 on either side of the shuffled log odds distribution). This procedure yields a proportion of events that significantly decode to the left (food) trajectory 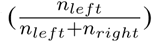 for each session, such that a score of 0.5 indicates an equal number of left and right events for that session, 0 indicates all right (water) events and 1 indicates all left (food) events. Consistent with the earlier analysis, food restriction sessions contained a larger proportion of significant events on the right (water) trajectory (left/food .25 ± .13, right/water .75 ±.13), and water restriction sessions shifted these proportions *away from* the right/water trajectory (left/food .52, right/water .48, difference *z* = –2.10, *p* = .035; Figure 2c). Again, each individual subject showed a negative difference score (Figure 2c), indicating that motivational state changes consistently shift significant SWR events away from the preferred outcome.

An important aspect to appreciate about this analysis is that a number of factors can result in a decoding bias for the left or right maze arm. These factors may be unrelated to the key experimental manipulation (motivational state), such as the number of cells with place fields on left vs. right trials, or idiosyncratic subject-specific behavioral experiences on one arm or another. Similarly, a “true zero” point corresponding to equal food and water motivation, if such an indifference point could be found at all, would necessarily be a highly transient state. Thus, we base our test of the hypothesis that motivational state affects SWR content on the *difference* between motivational states. Specifically, a given rat may show an overall SWR content bias for the right arm across both food- and water-restriction sessions, but the key question is what (if any) the difference in that bias is between motivational states. For instance, in the analysis of significant SWR proportions (Figure 2c), rats R042 and R064 had an overall SWR content bias toward the right (water) trajectory: the mean of the shuffled distribution (gray bars) is offset towards the right (likely related to differences in decoding accuracy; see Figure 6a). Crucially however, compared to this shuffled distribution, motivational state shifted SWR proportions significantly away from the preferred outcome. Phrased differently, for these two rats, the overall bias towards the right (water) trajectory was more pronounced during food restriction sessions compared to water restriction sessions. Importantly, this idiosyncratic bias is not required for motivational state to shift SWR content away from the preferred outcome: rats R044 and R050 do not appear to have an overall bias for the water (right) trajectory, yet show the motivational state shift.

The analysis so far included *all* SWR events, regardless of when in a session they occurred. However, the observed shift away from the preferred outcome may emerge within a recording session as a result of experience on the task that day (recall that all animals had previous experience with the task when neural recording began, see Figure 1a and *Online Methods*). Alternatively, SWR content shifts may be a direct result of motivational state changes alone and therefore *precede* experience that day. To distinguish between these scenarios, we computed SWR content separately for different task epochs. Confirming the previous analysis, an overall shift away from the preferred outcome was evident throughout task epochs, as can be seen from the consistently negative difference scores (*fr* - *wr*) in both left vs. right log odds (Figure 3a) and the negative differences in the proportions of significant left trajectory events (Figure 3b). In particular, ‘pre’ SWR food-vs. water restriction session log odds z-scores were –.50 ± .25 vs. .39 ± .32, difference *z* = –2.79, *p* =.005 based on resampled distribution; the ‘post’ difference was *z* –2.41, *p* = .016; Figure 3a). A similar pattern was present when proportions of significant events alone were considered (Figure 3b), although the pre-task proportions just failed to clear the conventional significance threshold (food restriction: proportion left (food) events .24 ± .13, water restriction: .53 ± .15, difference *z* = –1.93, *p* = .053). The post-task proportions differed significantly between food- and water-restriction sessions (*fr* .23 ± .12, *wr* .50 ± .13, difference *z* = –2.16, *p* = 0.032). Note that as before, each individual subject showed negative difference scores (Figure 3, insets), indicating a consistent bias in SWR content away from the preferred outcome. Computing log odds and significant events based on the decoded posterior past the choice point (i.e. maze arms only, rather than full trajectories; Figure S5) enhanced this effect (‘pre’ SWR log odds difference *z* = –3.11, *p* = .0019, ‘post’ *z* = –2.80, *p* = .0052; significant events only ‘pre’ *z* = –2.60, *p* = .0092, ‘post’ *z* = –2.85, *p* = .0044).

**Figure 3:**
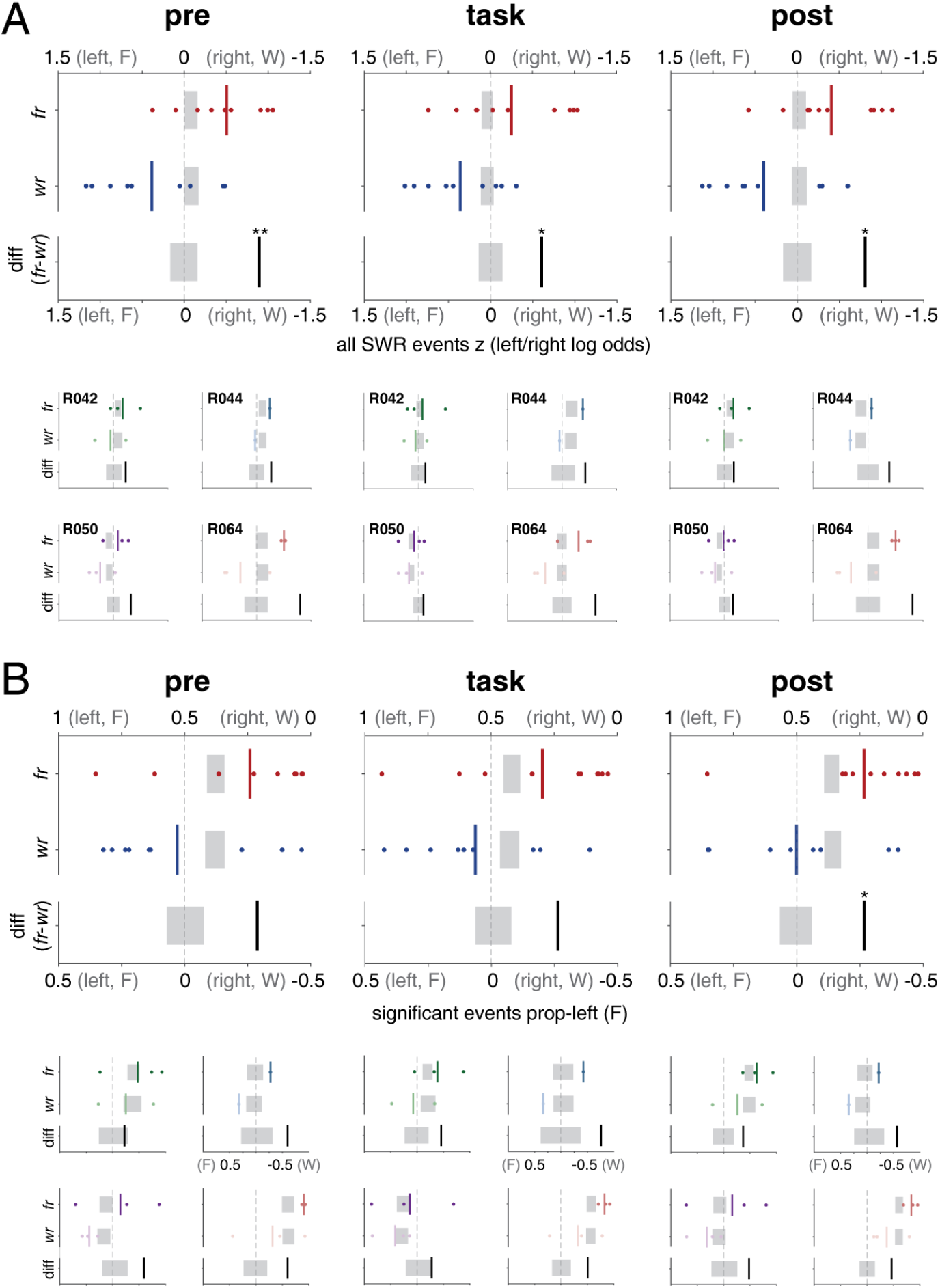
Sequenceless SWR content is biased away from the preferred outcome before experience on the task that day. **A**: Mean z-scored log odds (left vs. right, as in Figure 2b) across food-restriction sessions (*fr*, red), water-restriction sessions (*wr*, blue; upper horizontal axis), and their difference (*diff*, black; lower horizontal axis), plotted separately for each within-session epoch: before experience on the task that day (“pre”, left column), during intertrial intervals (“task”, middle column) and following experience (“post”, right column). **B**: Proportion of significance-thresholded SWR events that decode to the left (food) trajectory (as in Figure 2c) for food-restriction sessions (red), water-restriction sessions (blue; upper horizontal axis, and the difference between them (food restriction minus water restriction session proportions; lower horizontal axis), separated according to within-session epoch. Small panels show single-subject data; axis labels are omitted when identical to those in the average plots.

If experience on the task drives SWR content, there should be a SWR content difference between the ‘pre’ and ‘post’ epochs (see the schematic in Figure S9a). However, there was no evidence for such a change in either the SWR content log odds (observed ‘pre’ vs. ‘post’ difference –.09, expected value based on shuffling ‘pre’ and ‘post’ labels .00 ± .11, *p* = .44) or significant SWR proportions (difference –.02, shuffled mean .00 ± .05, *p* = .73). Because the lack of a significant difference does not constitute evidence for the null, we employed a two one-sided test (TOST) to formally determine equivalence ^14^. Specifically, we set a priori equivalent bounds based on the observed difference in pre-task SWR content between food- and water-restriction days (a difference of .89 ± .30 for log odds; .29 ± .14 for proportions). Then, we tested whether the observed difference between pre- and post-task rest SWR content (–.09 ± .11 for log odds, –.02 ± .05 for proportions) was significantly within the equivalent bounds (*t*_(22.75)_ = –10.91, p *<* .001 for log odds, *t*_(22.52)_ = –7.92, p *<* .001 for proportions; Welch’s t-test). Thus, we can rule out the possibility that SWR content changed between the pre- and post-task epochs to the same extent as the change between motivational states. In particular, strongly biased behavioral experience (e.g. towards the left (food) arm on a food restriction day) is not sufficient to overcome the motivational state SWR content shift away from the preferred outcome. Similarly, if SWR content on this task reflects planning towards the next goal, pre-task SWR content would be expected to be shifted towards, rather than away from, the preferred outcome (Figure S9b).

### Sequence detection

A strength of the above “sequenceless” decoding analysis is that it contains no free parameters beyond how SWR event candidates are detected. However, it cannot determine whether SWR event content is organized into a sequence of locations, or the properties of that sequence (forward, backward). Thus, we performed a complementary sequence-based analysis that detects sequences in the full decoded posterior according to a “maximum-jump” criterion and overlap with SWR event candidates ^4^ (*Online Methods*; see Figure 4 for example sequences). Analogous to the significant event detection procedure used earlier, we determined the proportion of significant left (food) sequences 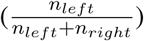 for each session, such that a score of 0.5 indicates an equal number of left and right sequences for that session, 0 indicates all right (water) sequences and 1 indicates all left (food) sequences. We plotted this proportion across the 6 alternating food/water restriction days (Figure 5a; see Figure S6 for raw sequence counts, and Figure S7 for single-subject data). As in the sequenceless analysis above, SWR sequence content was shifted *away from* the animals’ behavioral choices. On food-restricted days (red-shaded areas in Figure 5a), when animals preferred food (black line above 0.5), sequence content was more biased toward water (green line below 0.5) than on water-restricted days (blue-shaded areas), when animals preferentially chose water (black line below 0.5). A linear mixed model with subject-specific intercepts confirmed that motivational state predicted sequence content, changing the mean proportion of left (food) arm sequences from .35 for food-restriction sessions to .57 for water-restriction sessions (likelihood ratio test for models with and without motivational state: *χ*^2^ = 4.27, *p* = .038). Sequence content was anti-correlated with behavioral choice across sessions (*r*_(*n*=19)_ = –0.56, *p* = .014). Thus, motivational state changes shifted SWR sequence content in the opposite direction from the behaviorally preferred choice.

**Figure 4:**
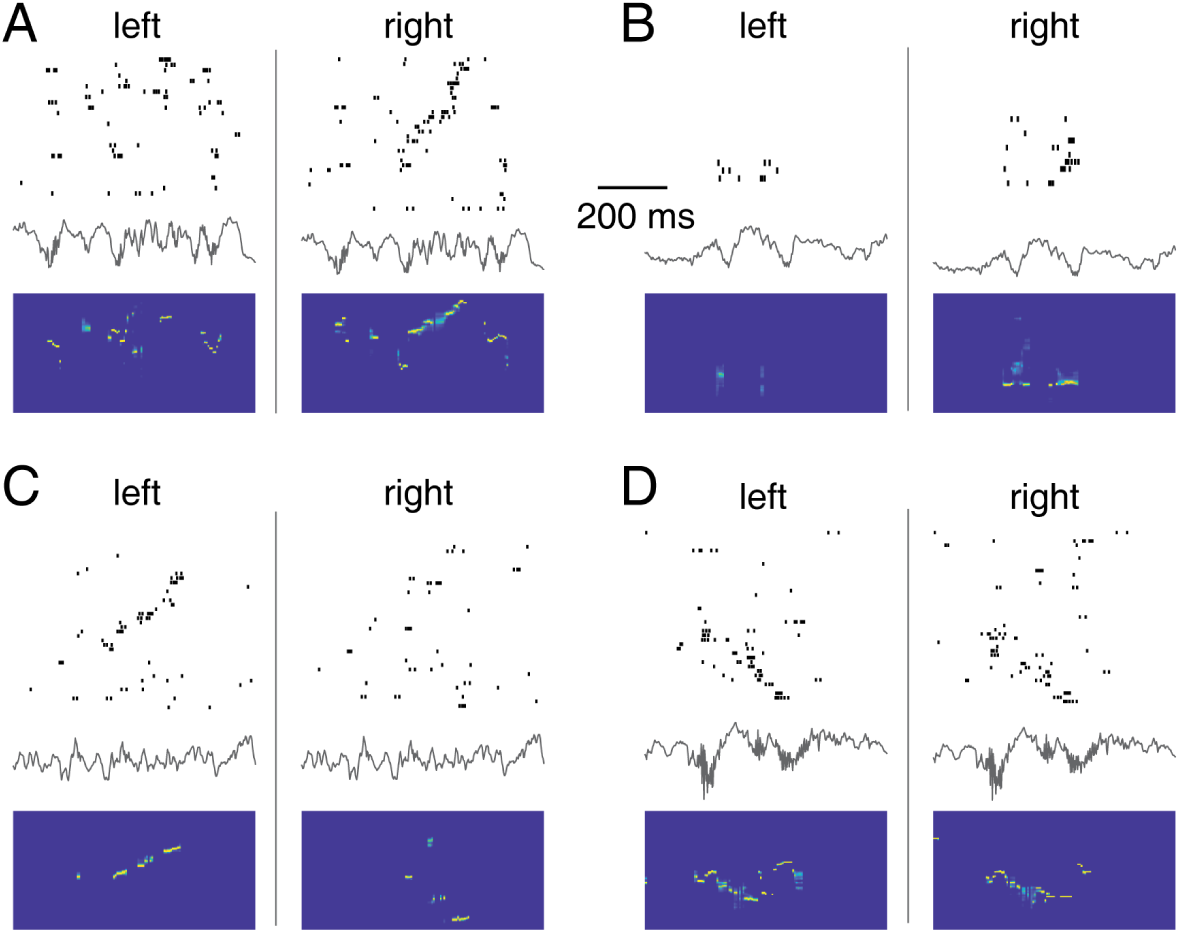
Hippocampal sequences detected during example sharp wave-ripple (SWR) events and their content. Panels **A-D** contain two columns, depicting spiking activity from the same time interval but using tuning curves obtained from “left” and “right” trials respectively. Within each column, a rasterplot (top) contains rows with tickmarks indicating spikes from a single neuron, ordered according to the location of their tuning curve peak firing rate on the track. Below the rasterplot is a local field potential trace (filtered between 1-425 Hz) and the decoded posterior probability distribution (blue: low probability, yellow: high probability). Each panel is an example from a different subject; examples were chosen from representative sessions and include a session with low (B, 53 cells total) and high (D, 162 cells total, note that only cells with place field(s) are shown in the rasterplots) cell counts. The events in panels A-C met the sequence detection criterion for only the right, right and left trajectory respectively and were scored as such; in contrast, the event in panel D met the sequence detection criterion for both left and right trajectories, and was therefore excluded from further analysis. In general, we found forward (A, C), reverse (D) and static sequences (B), which were considered both pooled and separately in further analyses.

**Figure 5:**
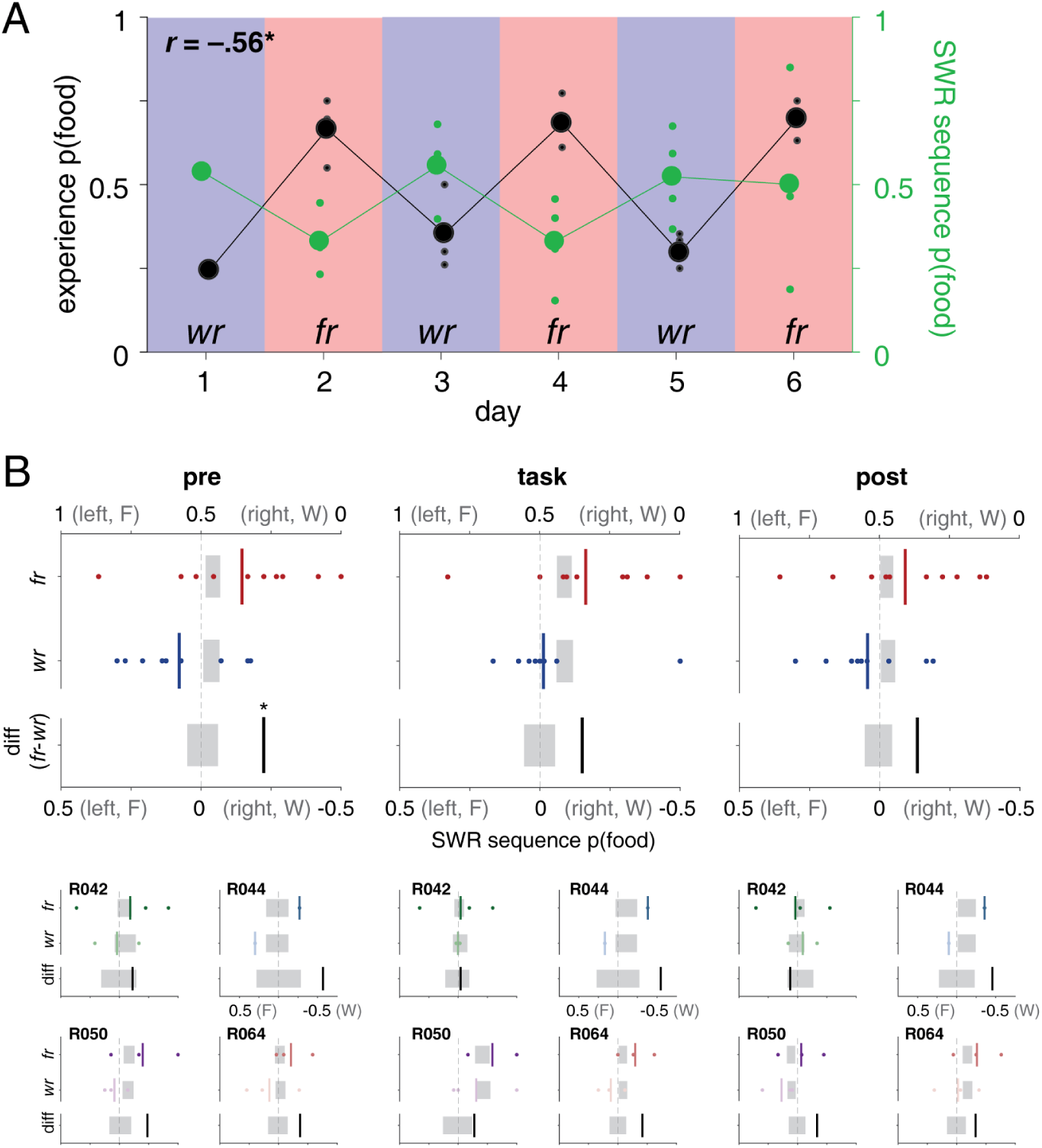
Motivational state changes shift SWR sequence content away from the behaviorally preferred maze arm. **A**: The proportion of detected SWR sequences that decode to the food (left) arm (green data points) is anticorrelated with behavioral preference (black data points). As in Figure 2a, blue shaded areas indicate water-restriction sessions and red shaded areas indicate food restriction sessions. **B**: SWR sequence content averaged across sessions is consistently biased away from the behaviorally preferred maze arm across task epochs (pre-task, task, and post-task); figure layout as in Figure 3.

As with the sequenceless results above, we next computed separately the proportions of left and right SWR sequences for different task epochs (Figure 5b). Confirming the previous analysis, an overall sequence content change opposite the motivational shift was evident, as can be seen from the consistently larger proportion of right (water) sequences on food-restriction days, which is shifted toward the left (food) on water-restriction days. Again, this shift was clearly present during the pre-task epoch, before experience on the task that day (Figure 5b, left panel; p(left) for food vs. water restriction, .35 ± .13 vs. .58 ± .09, difference *z* = –1.99, *p* = .046 based on resampled distribution). As in the sequenceless SWR content analysis (Figure 3), the observed difference between the pre- and post-task epochs was significantly within equivalent bounds defined by the pre-task food-vs. water-restriction difference (observed pre vs. post difference –.09, shuffled mean .00 ± .09, *p* = .32; two one-sided test *t*_(34.64)_ = –4.29, p *<* .001). As in Figure 3, there are some epochs and subjects in which the shuffled distributions are shifted to the right (water) arm, indicating an overall bias across motivational states to represent this arm (e.g. R064). Crucially however, the right (water) preference is larger for food than water restriction sessions, indicating that motivational shift consistently biased SWR sequences away from the preferred choice. This was generally the case for individual subjects and session epochs (Figure 5b, insets, note the single exception out of 12 non-overlapping analyses is R042’s “post” sequence content, which showed a small bias towards the preferred side). Thus, overall, the shift in SWR sequence content away from the preferred outcome preceded experience on a given day, and persisted throughout the session. (See Figure S8 for analyses on beyond-choice point sequences only, and forward vs. backward sequence comparisons, which show similar overall patterns.)

### Relationship with behavior

As shown above, decoded SWR content was anticorrelated with behavioral preference on the task on a session-by-session basis when using either the sequenceless or sequence-based measures. However, a possible explanation for this is that, rather than there being a direct relationship between SWR content and behavior, motivational state independently affects both. To distinguish between these possibilities, we performed a formal model comparison between linear mixed models (*Online Methods*). The inclusion of session-wide SWR sequence content improved the prediction of behavioral preference compared to motivational state alone (motivational state only, AIC 267.94; adding SWR content, AIC 259.06, likelihood ratio test *LRstat*_Δ*df* =1_ = 10.88, *p* < 0.001), suggesting that SWR sequence content bias and behavior are related beyond what is expected from independent effects of motivational state alone. To explore whether trial-by-trial changes in behavior were related to SWR sequence content, we constructed logistic regression models that sought to predict trial-by-trial behavioral choice based on a number of regressors, including (1) intertrial interval (ITI) content variables such as the proportion of left vs. right sequences, the most frequent and the most recent sequence, and (2) session-wide variables such as motivational state and session-wide replay content. Evidence of a true trial-by-trial relationship between SWR sequences and behavior requires that ITI regressors improve model fit relative to session-wide regressors alone. We failed to find any ITI regressors supporting such improvement, suggesting that either trial-by-trial differences in SWR content on this task were unrelated to behavioral choice, or there were insufficient ITI sequences and/or free-choice trials to enable a sufficiently powered test.

### Null hypothesis and scenarios

A crucial aspect of any analysis examining SWR content across different conditions, such as the left and right trials used here, is to compare the observed distribution against an expected distribution that takes decoding accuracy into account. For instance, imagine that for a given recording session, we happened to record many more place fields on the left arm than on the right arm; a strong SWR content bias for the left arm would then result from improved ability to detect significant SWR content, rather than a true underlying change in that content. Similarly, it is possible that subtle differences in the rats’ behavior on the preferred vs. non-preferred arm biased our ability to detect SWR content. The impact of these and other factors can be systematically quantified using a leave-one-trial-out procedure, which computes a cross-validated decoding error (when the animal is running) that can be converted into a null hypothesis of expected SWR content ^15^. The null hypothesis quantifies expected differences in *observed* SWR content that arise from limited and unequal sampling in the case that *true* underlying SWR content is equal between conditions.

Using this procedure, we found a small bias of lower error (higher accuracy) for behaviorally-preferred trials, containing the restricted outcome (Figure 6a), such that the expected null distribution of SWR sequence content favored the behaviorally-preferred trials (Figure 6b; this effect likely occurs because behavioral trials toward the preferred outcome were less variable ^15^). Note that this non-uniform null distribution favors the hypothetical result that motivational state biases SWR content *toward* the preferred outcome: to the left (food) for food restriction (*fr*) and to the right (water) for *wr* sessions. Importantly, the actual data show a shift in the opposite direction from this predicted null distribution (compare Figure 6b with Figures 3 and 5b). Our analyses also controlled for another potential source of bias, the number of trials used to obtain tuning curves for decoding, by subsampling the number of behavioral trials to be equal for left and right trials (*Online Methods*).

**Figure 6:**
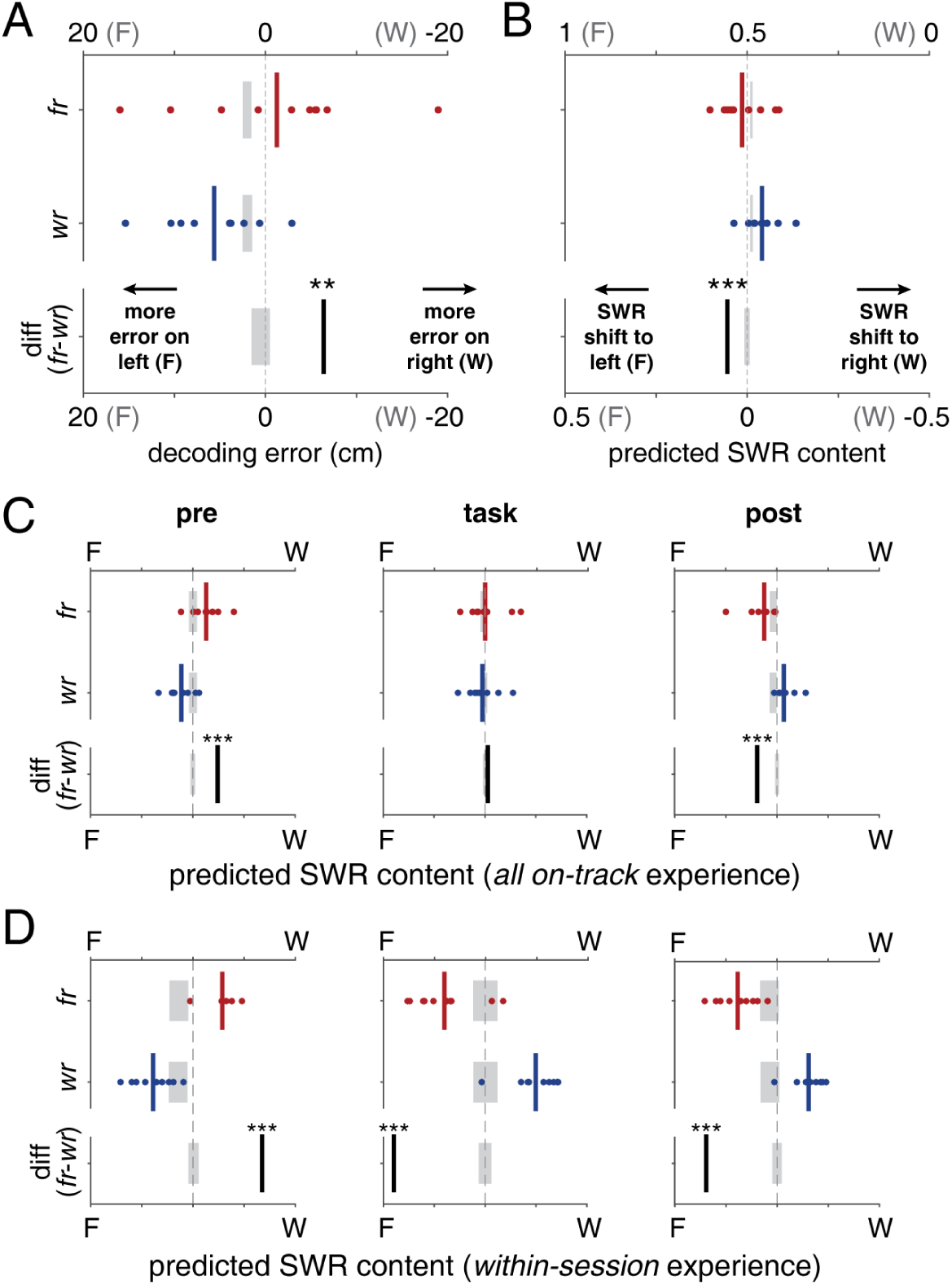
**A**: Decoding error during trials on the track, as obtained with a leave-one-trial-out cross-validation procedure ^15^. Each individual data point shows the difference between decoding error for left (food) trials and decoding error on right (water) trials for a single session. The decoding errors for food-restriction sessions (*fr*) and water-restriction sessions (*wr*) are shifted in opposite directions from the mean decoding error (center of the gray bar, which indicates the mean and SD obtained by randomly permuting session labels). Note that the mean decoding error across all sessions is larger on the left (food) arm. **B**: The difference in decoding errors between food- and water-restriction sessions in panel A leads to a predicted distribution of SWR content. This null hypothesis predicts a slight bias toward the arm containing the restricted outcome, whereas the data indicates the opposite (compare the left/food *diff* shift in this panel to the right/water shift in Figures 3b and 5b). Note that the shuffled distributions (gray bars) indicate a slight overall bias towards the right (water) arm. **C**: Expected SWR sequence content distributions according to a simulated scenario in which sequence content reflects accumulated experience across all recording days. **D**: As in C, except that sequence content reflects experience within single recording sessions, with pre-task content carried over from the previous day’s experience. Note that neither scenario correctly predicts the observed SWR sequence content (compare with Figures 3b and 5b).

Given that the observed sequence content cannot be explained by differences in the cross-validated decoding error, we next considered potential explanations based on subjects’ experience. Following Gupta et al. ^16^, we simulated scenarios in which each SWR candidate event in the data is probabilistically assigned a “left” or “right” label according to one of the following rules: (1) replay in proportion to experience accumulated over the six recording session days, (2) replay in proportion to within-session experience. For instance, say a SWR event was detected in the inter-trial period following trial 8 of day 2. If all trials that day were “left”, scenario (2) would assign “left” to that event. If all trials up to that point (including previous sessions) were 45% “right”, then scenario (1) would assign “left” to that event with 45% probability. Since in scenario (2) within-session experience has not occurred yet during the pre-task period, we carried over the post-task distribution from the previous day. The SWR sequence content proportions expected from these scenarios are shown in Figure 6c-d: although a pre-task bias similar to that in the data was present, the simulations incorrectly predicted a reversal of this bias with experience. Figure S9a visualizes these predictions alongside the pattern actually observed in the data, showing that the experience-dependent scenarios could not account for the observed data.

A further possibility is that SWR sequence content is experience-dependent, but with a delay (Figure S10a). Such a delay would be necessary to explain the observed opposite-side bias that persists in the data throughout the pre-task, task, and post-task epochs (Figures 3 and 5b; as the experience-dependent scenarios indicate, without a delay this bias would be expected to reverse between pre- and post-task epochs.) Experience-dependent increases in SWR frequency and content are thought to increase sharply immediately following experience, and then gradually decay ^17–19^. However, there are also observations of replay content that persist for hours and possibly days following experience ^13, 20^. Thus, we tested the hypothesis that behavioral bias on day *n – 1* should predict the pre-task SWR content bias on day *n*. (Note that behavioral bias, which varies between 0.5 (equally preferred) and 1 (all left, or all right) is different from raw behavior; see Figure S10.) For instance, a strong behavioral bias toward the food arm on day *n – 1* would result in a strong SWR content bias toward food during day *n*’s pre-task epoch (Figure S10a), opposite that day’s restriction type and behavioral preference (now switched to water). However, we found no evidence for correlations between behavioral bias on day *n – 1* and pre-task SWR content bias on day *n* (sequence decoding content bias, *r*_(17)_ = –0.11, *p* = 0.65, sequenceless decoding content bias, *r*_(17)_ = 0.26, *p* = 0.31). In contrast, and as would be expected from an account in which motivational state determines SWR content, pre-task SWR content bias predicted behavioral bias on the same day (*r*_(17)_ = .50, *p* = .028 for sequenceless SWR content; Figure S10b). Although sequence decoding SWR content bias did not yield a significant correlation, recall that the previously reported regression analysis shows that session-wide SWR sequence content improves the prediction of behavioral choice (section “Relationship with behavior”). Repeating this analysis with only *pre-task* SWR sequence content likewise resulted in improved prediction of behavioral choice compared to motivational state alone (likelihood ratio test, *LRstat*_Δ*df* =1_ = 6.71, *p* = 0.0096). Finally, motivational state on day *n* was a numerically better predictor of day *n* SWR content than day *n – 1* behavior (sequence content: behavior AIC 6.15, motiv. state 2.79; non-sequence content: behavior AIC 28.4, motiv state 27.67; lower values are better). Thus, although our experiment was not explicitly designed to rule out the hypothesis that experience on day *n – 1* drives replay content on day *n*, the best statistical account of the data is that motivational state biases SWR sequence content away from the preferred outcome.

## Discussion

*How do these results inform theories of the content and role of SWR sequences?* Two common views of the factors that determine SWR content are that (1) SWRs replay a recency-weighted record of experience, either as part of a dual-store model of memory ^21^ (usually attributed to SWRs during sleep) or as part of a retrieval process that can support ongoing decisions ^22^, both “retrospective”; and (2) SWRs depict trajectories toward possible goals, as part of a planning process ^4, 16^ (“prospective”). Our results are not well described by either of these accounts: simulations show that retrospective biases do not fit the data (Figure 6c-d and associated analyses; see Figures S9a and S10 a for schematic illustrations). Prospectively simulating a trajectory toward the non-preferred outcome as part of a planning process would certainly be informative in decision making ^23^. However, planning content is generally thought of as either agnostically considering options until some termination criterion, and/or containing a bias toward the goal (as in e.g. backward or goal-constrained search) ^3, 24, 25^. These prospective accounts would predict either a bias toward the chosen option, or no bias (see Figure S9b for a schematic illustration). Instead, we observed an opposite bias, reminiscent of Gupta et al. ^16^ who found that during left or right-only blocks on a T-maze, the opposite side was replayed more often than during alternation blocks. Here, we extend that observation of replaying the “other” option to include interoceptive cues (motivational shifts) that precede actual experience as a source of such non-temporal SWR content biases.

*If replay content on this task is not consistent with a memory trace of recent experience, or with prospective generation of upcoming choice, what could its function be?* There is an emerging appreciation that hippocampally-driven consolidation may involve processes beyond the simple offline training of cortical representations based on replay of time-limited information stored in the hippocampus. This broader concept of consolidation includes processes which would benefit from replay content that is not limited to recent experience ^26, 27^. Examples include the learning and maintenance of predictive “successor representations” ^28, 29^, and updating of decision variables in reinforcement learning (e.g. Dyna ^30^) which may prioritize states with high uncertainty ^31, 32^. When applied to the task used here, we speculate that hippocampal replay enables evaluation and updating of information related to the option not (about to be) taken. These ideas echo the less specific but related notion that replay could serve to prevent destructive interference of memory traces ^16^; alternatively, under certain conditions SWRs may be part of a forgetting mechanism ^33, 34^.

In any case, these possibilities would require that in the task used here, the pattern of SWRs observed reflects a hippocampal mode not necessarily engaged in supporting immediate behavior on the task, but operating in the service of longer-timescale behavior. This conceptualization blurs the distinction between the traditional online/offline modes to more of a continuum from highly task-constrained to more internally driven ^27, 35^. In this view, it is possible that even on T-maze tasks, highly overtrained animals may not be engaging their hippocampus (including SWRs) to perform the task, freeing it up for (re)consolidation processes. Consistent with this view, Singer et al. ^36^ found that the animals’ choice was predictable from SWR content only during a specific time window during learning, and the animals in Gupta et al. ^16^ had been extensively trained on the task prior to recording. Thus, an important limitation of these and many other hippocampal sequence recording studies is that it is often unclear to what extent SWRs support behavior on the task; even if hippocampal lesions or inactivations result in an overall impairment, this need not imply that each SWR directly supports performance. Regardless of the role of the hippocampus and SWRs on the present task, the observation that SWR content was modulated by motivational state highlights the nontrivial ways in which SWR content can relate to experience and reward revaluation in particular.

## Online Methods

### Subjects and timeline

Male Long-Evans rats (n = 4, Harlan; 4-8 months old when behavioral training began) were trained on a T-maze task, described in detail below, in daily recording sessions until they were running proficiently (range: 4-15 sessions). Subjects were surgically implanted with recording electrodes targeting the dorsal CA1 area of the hippocampus. Following recovery, subjects were re-trained on the task while electrodes were lowered to the CA1 cell layer and subjects were accustomed to running the task while plugged in to the recording system. Upon acquisition of a sufficiently large ensemble of units (range: 6-9 days from start of retraining), a 6-day sequence of daily recording sessions began, which were analyzed here. Note that this is the same data set used in van der Meer et al. ^15^ where we established methodological points referenced throughout the analyses reported here. All procedures were approved by the University of Waterloo Animal Care Committee (AUPP 11-06). Animals were kept on a regular day/night cycle (daylight from 7 am to 7 pm) and were run during the light phase.

### Surgery and recording probes

Electrode arrays were assembled from a custom 3D-printed base (Shapeways), machined metal parts (00-80 threaded rods and stainless steel cannulas, Component Supply Company; custom fabricated metal nuts and shuttles), and an electrode interface board (EIB-36-16TT, Neuralynx) to contain 15 or 16 independently movable tetrodes (Sandvik 12.7 *µ*m insulated NiCr wire, gold-plated to 300-400 kOhm impedance measured at 1 kHz) and 1-4 reference electrodes. Surgical procedures were as reported previously ^37^. Briefly, animals were anesthetized with isoflurane, the dorsal surface of their skull was exposed, and a ground screw was placed through the contralateral parietal bone. Electrode arrays were lowered to the surface of the cortex, and the remaining exposed opening was sealed with a silicone polymer (KwikSil). Then, the arrays were anchored to the skull using small screws and acrylic cement. Rats were given a minimum recovery period of four days, during which antibiotics and analgesics were administered, before retraining began.

### Behavioral procedures and apparatus

The apparatus was an elevated T-maze, constructed from wood, painted matte black with white stripes applied to the left arm (Figure 1a) and placed on a metal frame. Rats were trained to run trials on the elevated T-maze in daily sessions, having either been food or water restricted overnight (16 hours). Before beginning the task each day, rats were first given a small quantity of food (90 mg, 2 pellets) and 6% sucrose water (0.1 ml) in their home cage. (This was done because the instrumental conditioning literature shows that such re-exposure following revaluation is required for behavioral change to manifest during a subsequent extinction text, a phenomenon known as incentive learning ^38^.) Next, they were placed on a movable platform (a terracotta flower pot lined with towels) for a 5-20 minute pre-task rest period. Rats were trained over a period of at least 4 days to make a left or right turn at the choice point for food (five TestDiet 5TUL pellets, 45 mg each, dispensed using an automated feeder; Coulbourn) or 0.13 ml 6% sucrose water (dispensed using tubing equipped with an electrically controlled valve) respectively. Food and water locations remained constant throughout all training and recording sessions used here. A handheld barrier was used to prevent movement in the reverse direction. Rats were allowed to remain at the end of the goal arm until the reward was consumed, or until it was clear that the rat was not interested in the reward, before being transferred to the nearest pedestal where they remained until the beginning of the next trial. Each intertrial interval (on the platform) began at a length of two minutes, and decreased with each trial to as short as one minute or less. A stationary barrier was placed at the choice point during some trials in order to force the rat down the unfavored arm a minimum of five times per session so that place fields could be estimated for both sides of the track. These blocked trials were not included in behavioral choice data (Figure 1b-c) but were included in behavioral experience data (Figures 2a and 5a and associated analyses). Training occurred in a fully lit room during the light phase. Recording sessions proceeded in the same way as training sessions, with minor modifications lengthening the pre-task period to 20-30 minutes, followed by 40 minutes of the task, and finally 5-15 minutes of post-task recording on the pedestal. Following completion of daily recording sessions, rats were given access to food and water for *∼*4-6 hours, consuming large amounts of the outcome that was previously restricted, indicating that they had not reached satiety from rewards consumed during the task.

### Data acquisition and preprocessing

Neural activity from all tetrodes and references was recorded on a Neuralynx Digital Lynx SX data acquisition system equipped with HS-36 headstages and a motorized commutator. Local field potentials were continuously sampled at 2 kHz, and recorded relative to a reference located in the corpus callosum. Spike waveforms were sampled at 32 kHz for 1 ms when the voltage exceeded an experimenter-set threshold (typically 50 *µ*V) and sorted offline (KlustaKwik, K. D. Harris and MClust 3.5, A. D. Redish). Highly unstable cells were excluded from analysis, as well as cells with fewer than 100 spikes over the entire recording session, or poor isolation statistics. A video tracking system recorded the rat’s position based on headstage LEDs picked up by an overhead camera. Signals were preprocessed to remove chewing artifacts and high-amplitude noise transients where necessary.

### Session inclusion criteria

As in van der Meer et al. ^15^, recording sessions were included for analysis if at least 40 cells total were recorded. This left 19 of 24 total sessions (6 each from R050 and R064, 5 from R042, and 2 from R044).

### Data analysis overview

All analyses were performed using MATLAB 2017a and can be reproduced using code available on our public GitHub repository (http://github.com/vandermeerlab/papers). Data files including metadata are publicly available on DataLad (http://datasets.datalad.org, ‘MotivationalT’ data set). We first provide an overview of the major data analysis steps, and then discuss each step in more detail.

- **Estimating encoding models (tuning curves).** From the behavioral data on the task (position tracking) and spiking data, we estimate the relationship between location on the track and unit firing rates. We estimate these tuning curves in three conceptually and procedurally distinct sets. The first set separates left and right trials and is used to evaluate decoding accuracy when animals are running on the task using leave-one-out cross-validation ^15^. A similar set of tuning curves, without leave-one-out, is used for the sequence-based decoding analysis. The third set concatenates left and right trials in a single tuning curve and is used for sequenceless decoding. A key step used for all sets is equalizing the number of trials used for “left” and “right” tuning curves to prevent biases arising from unequal sampling.
- **Decoding.** Using the above tuning curves, we decode (1) running data for evaluating decoding accuracy to obtain a null hypothesis for decoded sequence content, and (2) SWR-associated activity only for sequenceless decoding, and (3) all activity for subsequent SWR-associated sequence detection. Key parameters of these steps include the kernel width used to obtain spike density functions, time step and window size, and the minimum number of cells active during each decoding window.
- **Sequenceless decoding content.** Because sequenceless decoding is based on tuning curves that include both left and right trials in parallel, the decoded posterior probabilities for both can be compared, and converted into a z-scored log odds relative to a shuffled distribution, reflecting SWR content.
- **Sequence candidate detection and selection.** Using the full-session decoded posterior, sequences are detected using the following key parameters: maximum jump distance, time gap allowed before sequence is broken, and minimum sequence length. A subsequent selection step only retains those sequences that meet certain criteria, such as coinciding with a SWR candidate and passing a shuffle comparison, as described below. Further analyses apply additional criteria such as only analyzing sequences that occurred during a specific task epoch, that have certain properties such as depicting a forward or reverse trajectory or specific part of the track, et cetera. Only sequences for either left or right trials, but not both, are retained.
- **SWR detection.** This step detects SWR events based on local field potential and multi-unit activity. In the sequenceless decoding analysis, these candidate SWR events are what is decoded (a single time window for each event). In the sequence-based decoding analysis, only candidate sequences that overlap with a SWR event are kept for further analysis.

#### Estimating encoding models (tuning curves)

Position data was first linearized by mapping position samples to the nearest location on experimenter-drawn idealized trajectories for left and right trials separately. Linearized trajectories were binned using approx. 3 cm per bin, such that the 334 cm track length resulted in 111 bins (except for one subject, R042, who used a track length of 257 cm with the same number of bins). Next, “run” epochs were identified when the rat’s speed (estimated using an adaptive filtering procedure) exceeded a fixed threshold (approx. 4 cm/s). A matched set of trials was identified such that (1) the number of left and right trials was equal, and (2) the time difference between the left and right trials was minimized. This matching procedure was necessary to prevent biases in decoding accuracy for left and right sequences due to differences in how tuning curves for left and right trials were estimated. Using these matched trials, raw average firing rates for left and right trials were estimated separately from run epochs only by dividing spike counts by occupancy for each linearized bin. Raw firing rates were smoothed with a *σ* = 1 bin kernel ^15^. Bins at the start and end of the track were excluded from analysis. As explained in the next section, tuning curves were estimated from different sets of trials on the track, as required for analysis of SWR sequences and analysis of decoding accuracy.

#### Decoding

We ran three separate decoding analyses. The first analysis decodes each SWR as a single time window and determines its content (sequenceless decoding). The second analysis decodes all the data and then runs a sequence detection algorithm, and the final analysis decodes only running data (not SWRs) to establish decoding accuracy for left and right trajectories (a control analysis which provides a null hypothesis for SWR content across conditions). These analyses differ in (a) which trials were used to estimate tuning curves, (b) what data was decoded, and (c) what further analyses were ran on the output of the decoder (all described in detail in the next sections). All decoding analyses used the standard memoryless Bayesian decoding procedure ^12, 15, 39^.

For *SWR sequenceless decoding*, we first constructed tuning curves for left and right trials as described above, and then concatenated them into a single [*nCells* x 2 *∗ nLocationBins*] matrix, keeping track of which locations corresponded to left and right trials respectively. Next, we used the putative SWR event intervals found by the SWR detector (described below) to decode each event as a single time bin. The decoded posterior was summed for all left and right bins separately and converted into a log odds 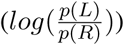. Repeating this procedure 1000 times for tuning curves in which random subsets of spatial locations had their left and right tuning curves swapped yielded a shuffled distribution which was used to obtain two SWR content measures: a z-score (averaged over all events in a session, yielding a single number) and a significant event count. The latter was obtained by including only those events that exceeded either the left- or rightmost 5% of the shuffled distribution, yielding left and right counts for each session. Only SWR events with at least 4 neurons active were included. To compare SWR content across multiple sessions, we simply computed the average difference in SWR content between food-restriction and water-restriction sessions. For example, for significant SWR events, this is [*p*(*L*)*_f r_ − p*(*L*)*_wr_*] where *p*(*L*) are the proportions of events depicting left and right trajectories on the track respectively, and the *fr* (food restriction) and *wr* (water restriction) subscripts indicate the session restriction type.

For *SWR sequence decoding*, we first obtained “left” and “right” tuning curves with matched trials as described above, and used these to obtain left and right decoded posteriors by decoding spiking data from the entire session. Spike trains were converted to spike density functions by convolving with a *σ* = 2 ms Gaussian kernel ^15^. We used the canonical one-step Bayesian decoding procedure with a uniform spatial prior and a 25 ms moving window, which was stepped in 5 ms increments. At least 4 cells had to be active in each window for decoded output to be produced. The output of this decoding procedure was then used for sequence detection analysis, described below.

For the analysis of decoding accuracy, we decoded each trial separately, using tuning curves obtained from all other trials of the same type (left or right). This leave-one-out procedure tests how well tuning curves generalize to the decoding of withheld trials, a situation that mimics some aspects of generalizing tuning curves to decoding of SWR sequences ^15^. The output from this decoding procedure was used to compute decoding error as the mean distance between the maximum likelihood (or equivalently, maximum a posteriori, since we used a uniform prior) decoded location and the true, observed location. Decoding parameters were the same as above (25 ms window, *σ* = 2 ms, minimum 4 cells active per bin). To derive the null hypothesis of sequence content, decoding errors were converted into proportions as 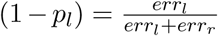 and 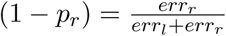.

#### Sequence detection and selection

To detect candidate sequences from the decoded output, we applied the procedure introduced by Pfeiffer and Foster ^4^. A candidate sequence starts at the first available sample and extends to the next sample, unless the distance between decoded locations (defined as the bin with the maximum likelihood) in two adjacent time steps exceeds a threshold (40 cm unless described otherwise) or no decoded location was available (due to insufficient number of cells being active at that time step). These conditions break the candidate sequence, and a new one is started at the next available decoded sample. The resulting pool of candidate sequences is then subjected to further criteria, starting with a minimum length (50 ms), a minimum number of neurons participating in the event (4), overlap with a putative SWR for any non-zero amount of time (see the section below for how these are detected). Next, for each time bin of each remaining candidate sequence, we computed in what fraction of identity-permuted shuffles (n = 500) that bin was also included in a sequence. Specifically, for each individual identity shuffle, we decode and detect sequences in the entire session as above (but now based on shuffled tuning curves). Across identity shuffles, we can then track how often any individual bin was labeled as being part of a sequence. Only observed candidate sequences in which the average proportion of bins in a sequence in the shuffles was 0.05 or less were retained for further analysis. Finally, because this analysis was performed separately for “left” and “right” trajectories, based on their respective tuning curves, the final step only retained those sequences that were classified as either left or right, but not both. More precisely, any ambiguous sequence that overlapped for any amount of time with any opposite-side sequence resulted in removal of both sequences.

The results reported in the main text used our default analysis parameters, which allowed no “jumps” or “skips” in the sequence detection and included all sequences, regardless of whether they were forward, reverse, or neither. In the supplementary figures we report results with different parameter settings and selection steps, including for sequences that are forward, reverse, and other restrictions. To determine if a sequence was forward or reverse, the beta coefficient for the time regressor needed to be significantly different from zero in predicting location in a linear regression (t-statistic, *p* < .05); because animals ran the track in one direction only, positive betas indicate forward sequences and negative betas indicate reverse sequences.

#### Sequence content analysis

The output of the above procedure is, for each session, a set of detected sequences, which are intervals (start and end times) with a label indicating which trajectory on the track they depict (left/right). Because the raw number of sequences detected will vary substantially due to factors of no relevance to the main questions of interest, such as the number of cells recorded or the number of SWRs detected on any given day, the main analyses use *proportions* of sequences. As above, used the fraction of sequences for each trajectory and the total number of sequences, e.g.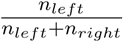 for left sequences. To compare sequence content across multiple sessions, we used [*p*(*L*)*_f r_ − p*(*L*)*_wr_*] as in the sequenceless analysis above.

#### SWR event detection

For the results in the main text, we used a thresholding procedure based on spectral content of the local field potential, combined with adaptive thresholding of multi-unit activity, to identify putative sharp wave-ripple complexes. Briefly, for each recording session, a noise-corrected SWR Fourier spectrum is obtained by manual identification of example SWR events, and subtracting a non-SWR “noise” Fourier spectrum. The dot product between this noise-corrected frequency spectrum and the frequency spectrum of the LFP is then obtained using a sliding 60-ms window to obtain a “SWR similarity score”. Separately, a multi-unit activity detector was defined, starting from a spike density function obtained by convolving the spiking activity of all recorded units with a *σ* = 20 ms kernel. An adaptive contrast enhancement step was applied to the raw MUA to emphasize transient changes in firing. Finally, a composite SWR detector was obtained by taking the geometric mean of the noise-corrected frequency spectrum and the modified MUA. The resulting unitless quantity was thresholded at 4 to yield candidate events, which were further restricted by the following requirements: (1) subjects could not be moving faster than 4 cm/s, (2) theta power could not be more than 2 SDs above the mean, (3) minimum duration of 20 ms, and (4) minimum 5 units active. We believe this method performs the best of those we have tested in identifying true SWR events while minimizing false positives. Rats appeared to be mostly awake during recording sessions, although it is likely some SWRs in the pre- and post-task rest periods occurred during early sleep stages.

#### Statistical analyses

All major analyses compare a dependent variable of interest across two different motivational states (food-restriction vs. water-restriction sessions). Thus, the effect size and statistical significance of motivational state on these variables of interest is given by the comparison of the true (observed) difference and a distribution of resampled values, derived from random permutations of motivational state labels across recording sessions. Throughout the paper, the mean and standard error across 1000 such “shuffles” are shown visually as gray bars in the figures, with statistics reported as *z*-scores and associated significance levels in the text. Note that in each shuffle, a session is assigned either a food- (*fr*) or water-restriction label (*wr*), but not both. Therefore, assigning only the *fr* labels fully specifies that the remaining sessions are *wr*, and vice versa. For this reason, the permutation z-scores are identical in magnitude when comparing *fr* sessions vs. shuffle, *wr* sessions vs. shuffle, and their difference (*diff*) vs. shuffle. Thus, single z-scores and associated p-values are reported in the text for each permutation test, visually indicated by asterisks above the *diff* data in the figures.

The effects of different behavioral variables on SWR content are tested with two further analyses. First, we fit linear mixed models with subject-specific intercepts to SWR content, and tested whether the addition of motivational state and/or behavior improved model fit (*fitglme()* function). Results reported are for best model fit as determined by the lowest AIC and BIC (which were in agreement across all comparisons), among alternative models fit with and without subject-specific intercepts. Formal model comparisons are performed with a likelihood ratio test (Williams, 2001; MATLAB *compare()* function). Additional regression analyses use this same overall approach, albeit with different regressors and dependent variables as indicated in the text. Second, we correlate SWR content with behavior across sessions (Pearson’s *r*) and associated *p*-values, computed with the MATLAB *corrcoef()* function.

## Acknowledgments

We thank Nancy Gibson, Martin Ryan and Jean Flanagan for animal care and Min-Ching Kuo, Julia Espinosa and Eric Carmichael for technical assistance. We thank Elyot Grant for developing the SWR detection method used for the main analyses in this paper. This work was supported by the University of Waterloo and Dartmouth College (start-up funds to MvdM), the Netherlands Organization for Scientific Research (NWO, grant 863.10.013 to MvdM), and the Human Frontiers Science Project (HFSP, grant RGY0088/2014).

## Author contributions

AAC performed experiments and pre-processed the data. AAC, YT and MvdM wrote analysis code. AAC and MvdM performed data analysis. MvdM wrote the paper with comments from AAC and YT.

## Supplementary Figures

**Figure S1:**
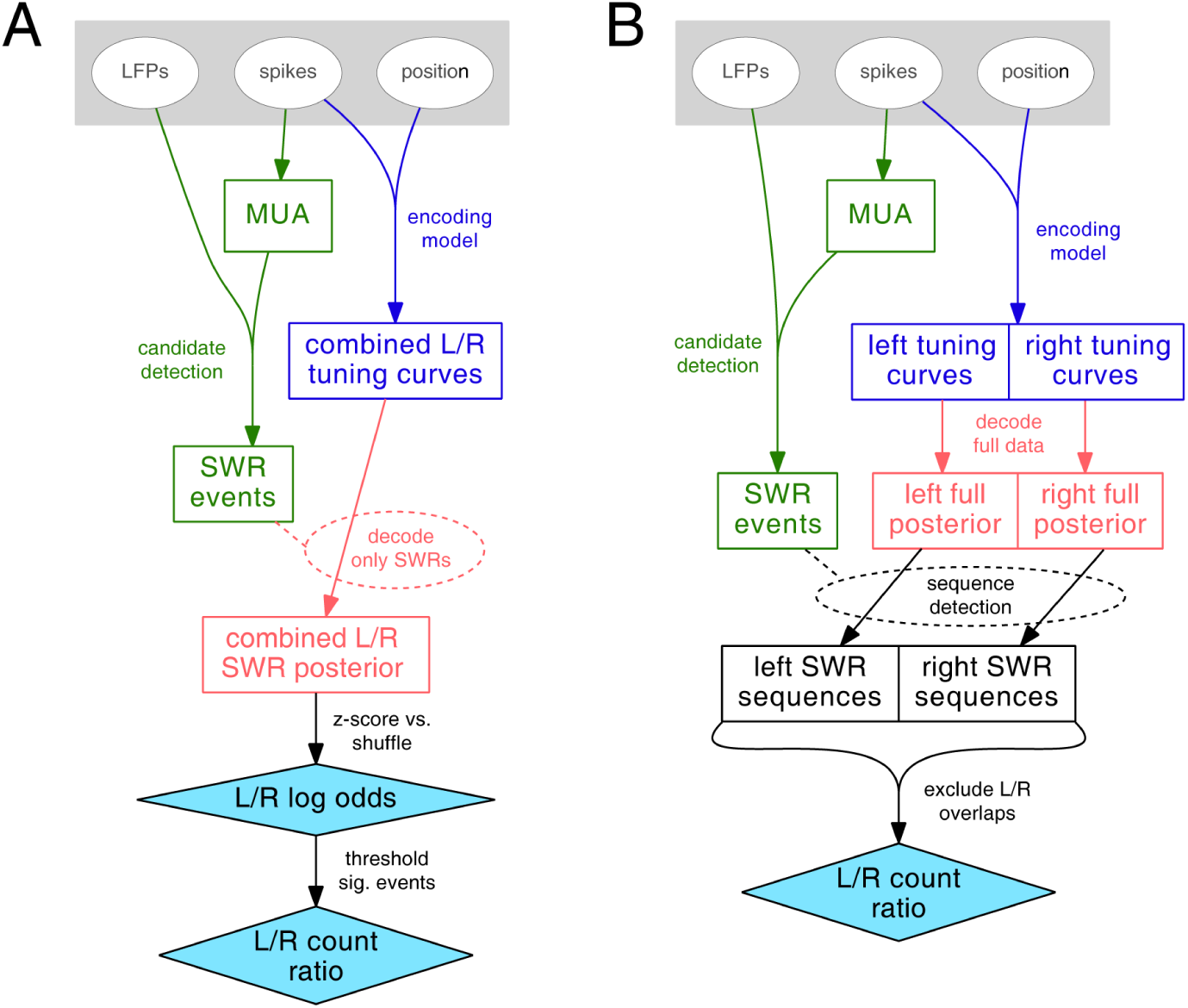
Schematic of SWR content analysis using sequenceless decoding (**A**) and sequence-based decoding (**B**). In the *sequenceless decoding* analysis, each candidate SWR event is decoded as a single time bin, using joint tuning curves containing left and right trial data. This analysis produces two outputs reported in the paper (drawn as cyan diamond shapes): the z-scored left vs. right log odds averaged across all events, and a left vs. right count ratio of significant events only. In contrast, *sequence-based decoding* (**B**) first decodes all data in a given session for left and right trial tuning curves separately, using a 25 ms moving window (step size, 5 ms). Next, sequences are detected in the left and right posteriors separately, before removing sequences that (1) did not overlap with a candidate SWR event, (2) were not sufficiently distinct from the number of sequences obtained from a random resampling procedure, or (3) were sequences for *both* left and right.

**Figure S2:**
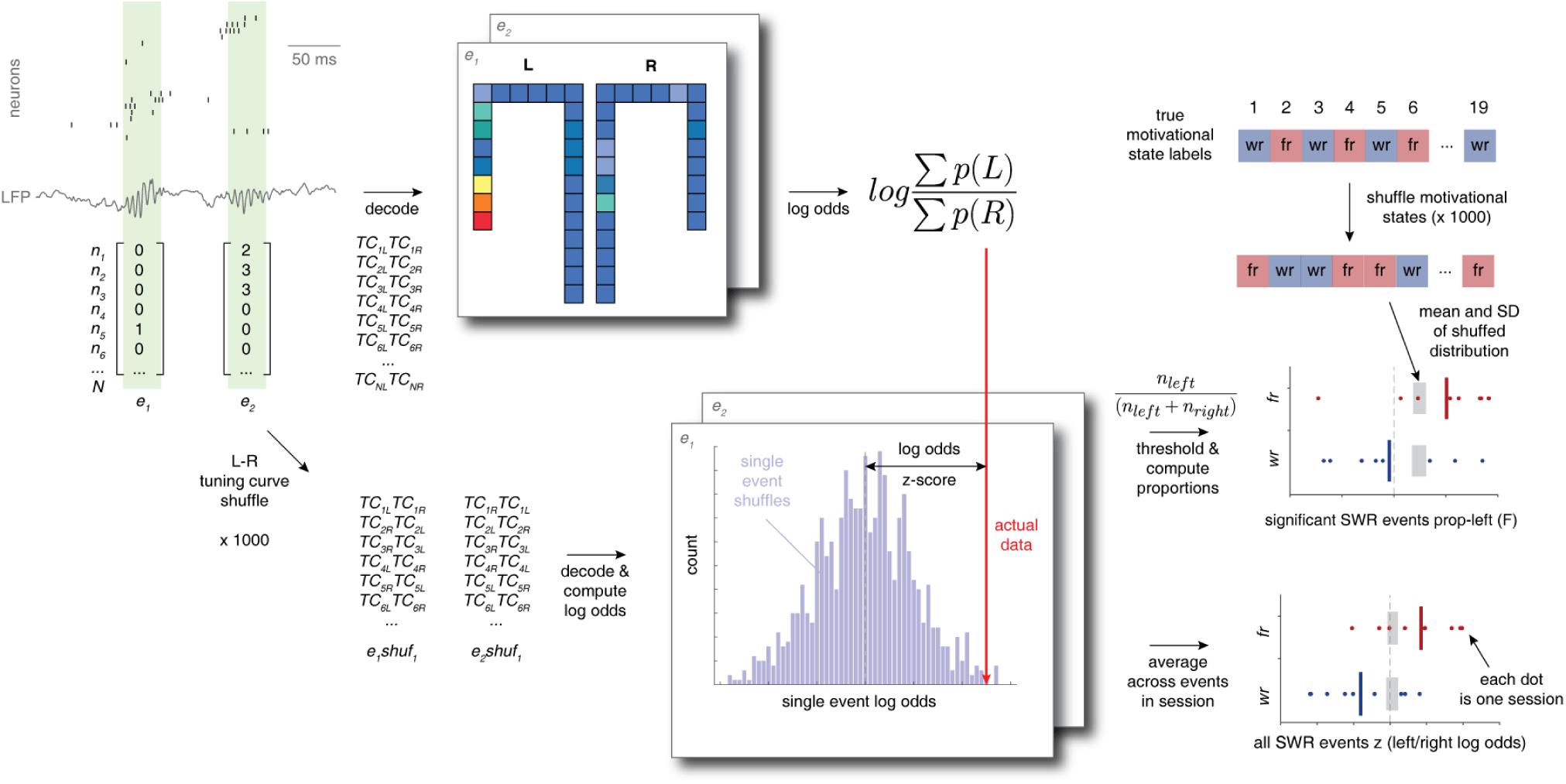
Schematic of sequenceless decoding analysis. Each SWR event (top left) is converted into a vector of spike counts and decoded into a joint probability distribution that includes both left and right trajectories of the T-maze. This probability distribution yields a left vs. right log odds score for each event. Because any left or right bias in this score may simply be due to unequal distributions of the number of place cells, average firing rates, and so on, this raw log odds score is compared to a distribution of log odds obtained from 1000 permutations of left and right trial tuning curves used in the decoding. This comparison yields a *log odds z-score* for each event, which is either averaged across all events in a session (bottom right figure and Figure 2b), or thresholded to keep only significant events to yield a proportion of left events (Figure 2c). To determine if SWR content is related to motivational state, both measures are averaged across food-restriction (*fr*) and water-restriction (*wr*) sessions, and the resulting values (black vertical bars in bottom right plot; dots indicate single sessions) compared to a distribution of averages obtained from randomly permuting food- and water-restriction labels across sessions (top right; gray bars indicate mean and SD of this shuffled distribution). Thus, this analysis uses two independent bootstraps: the first “tuning curve shuffle” quantifies SWR content on an event-by-event basis, and the second “motivational state shuffle” quantifies the effect of motivational state on SWR content averaged across sessions.

**Figure S3:**
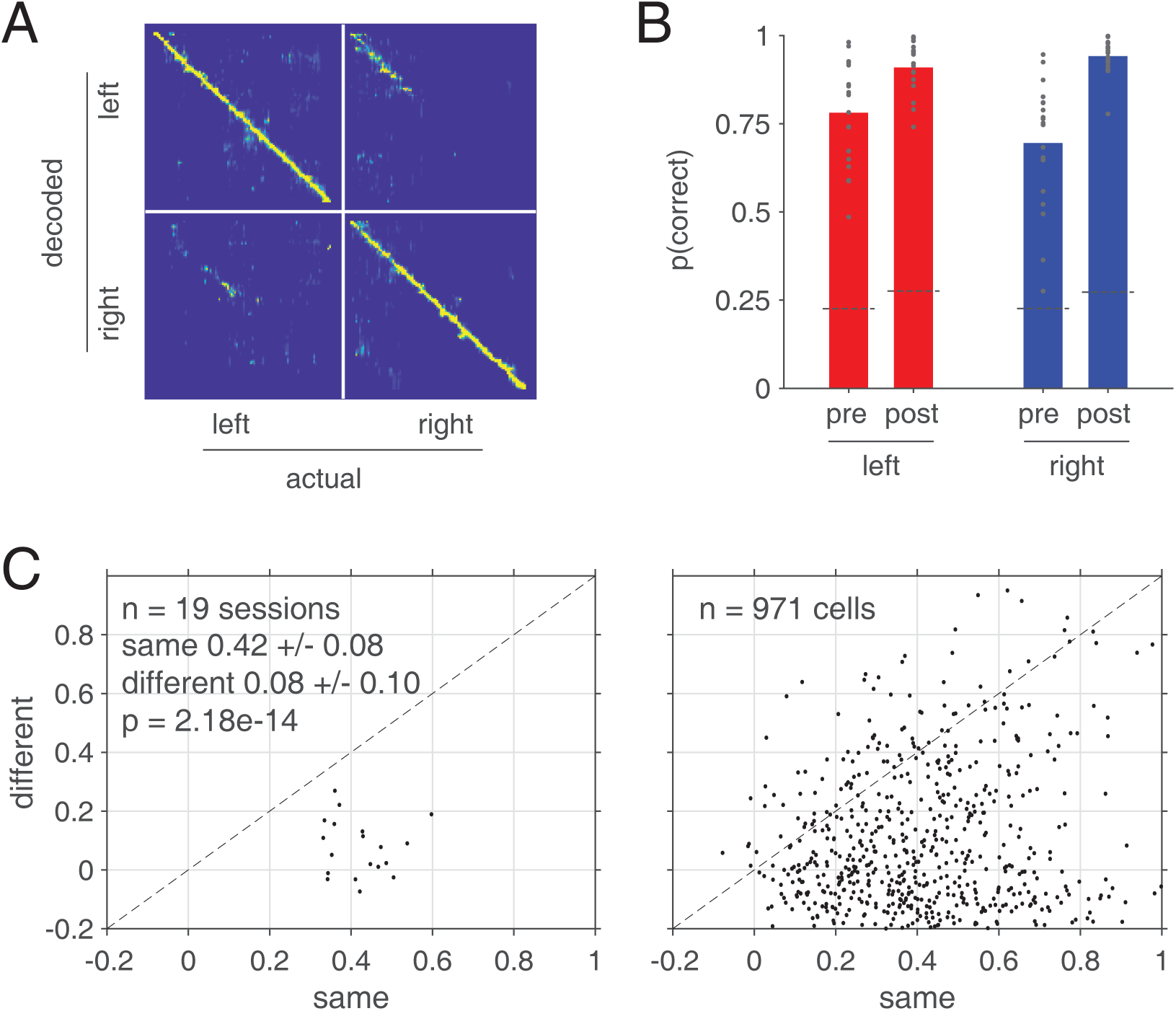
Left and right trials are clearly distinguishable during running on the track. **A**: Decoding confusion matrix for a single example session. Each decoded time bin (bin size = 100 ms) is assigned to the corresponding actual trial type (left, right) and position of the animal (horizontal axis). Decoded posteriors for all bins at that location are averaged to obtain the values in each column of the matrix. Perfect decoding would result in all diagonal elements being 1. Note the clear diagonal indicating non-random decoding overall, and the fainter, off-diagonal elements corresponding to confusion of left and right trials. **B**: Average classification performance across all subjects and sessions, based on the trial type (left, right) and location (before the choice point, ‘pre’; or after, ‘post’) of the maximum a posteriori decoding. Note that for all trial types and locations, classification performance is clearly above chance (all *p* < .001). Overall, classification performance was better for the right (water) arm compared to the left (food) arm (post-CP .92 ± .04 for right, .83 ± .06 for left, *p* = .009; *n.s.* for pre-CP). Importantly, there was no indication that classification performance differed between food- and water-restriction sessions (all comparisons *n.s.*). **C**: Single-trial tuning curve correlations between trials of the same type (left-left, right-right) are systematically higher than correlations between trials of different type (left-right), both when averaged across all cells within a session (left panel) and on a cell-by-cell basis (right panel). Only positions on the common, central stem of the maze were included in this analysis. The higher correlations between trials of the same type, compared to the correlations across types, indicate left and right trials are distinct. Thus, as measured by two different approaches, left and right trials are clearly distinguishable during behavior on the track, even on the common portion of the maze.

**Figure S4:**
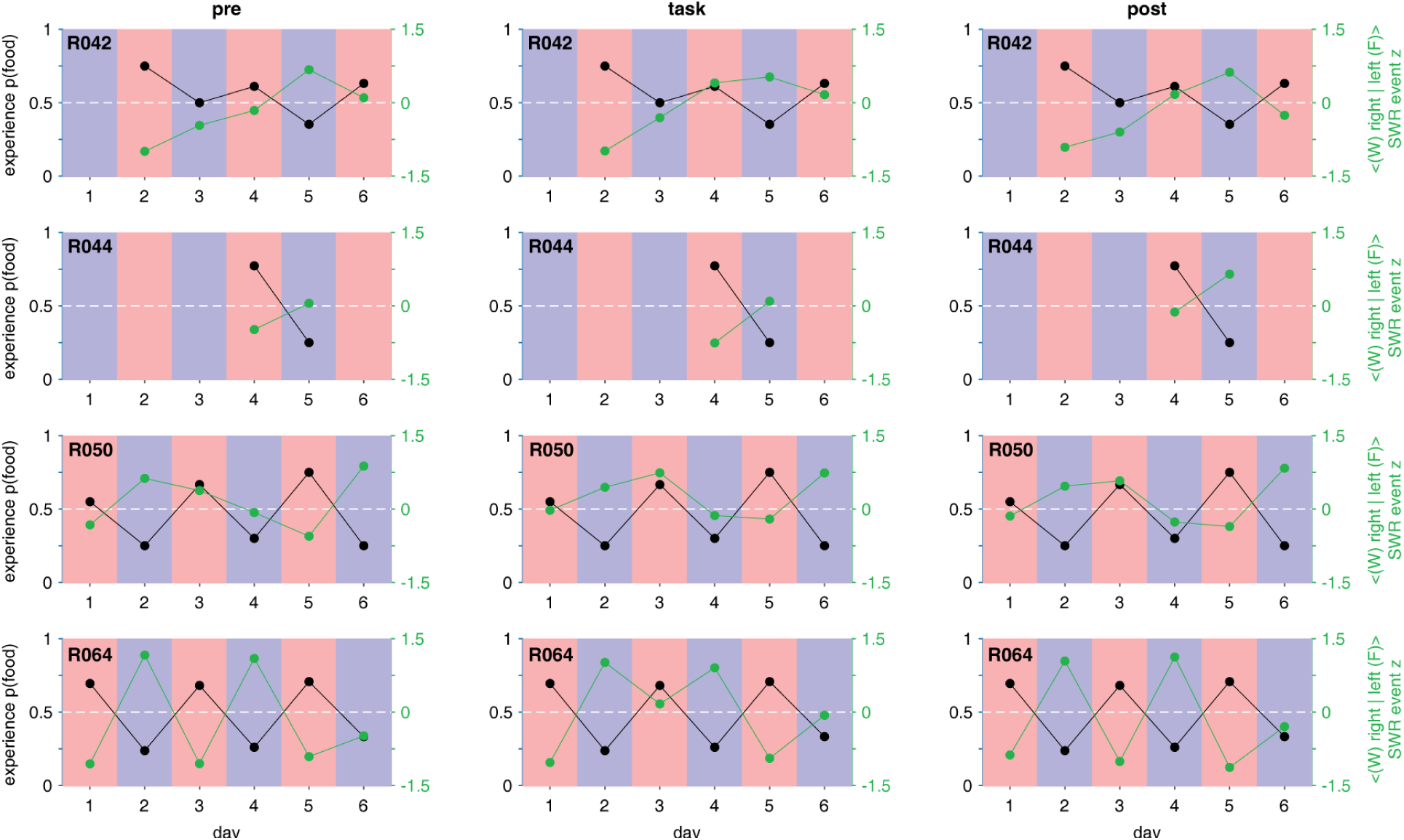
SWR content z-scored log odds for each individual epoch (columns correspond to pre-task rest, task, and post-task rest respectively), session (each data point corresponds to a single session), and subject (each row shows data from a single subject). Individual subjects may show idiosyncratic biases such as an overall shift in SWR content towards the water (right) arm, but in each subject and epoch *changes across days* in SWR content tended to be opposite changes in behavior. For instance, in the pre-task (left) column, it can be seen that this is the case for all pairs except for R042 day 3 to 4, and R050 day 3 to 4. In other words, 13 out of 15 motivational shifts resulted in a SWR content shift opposite from behavior.

**Figure S5:**
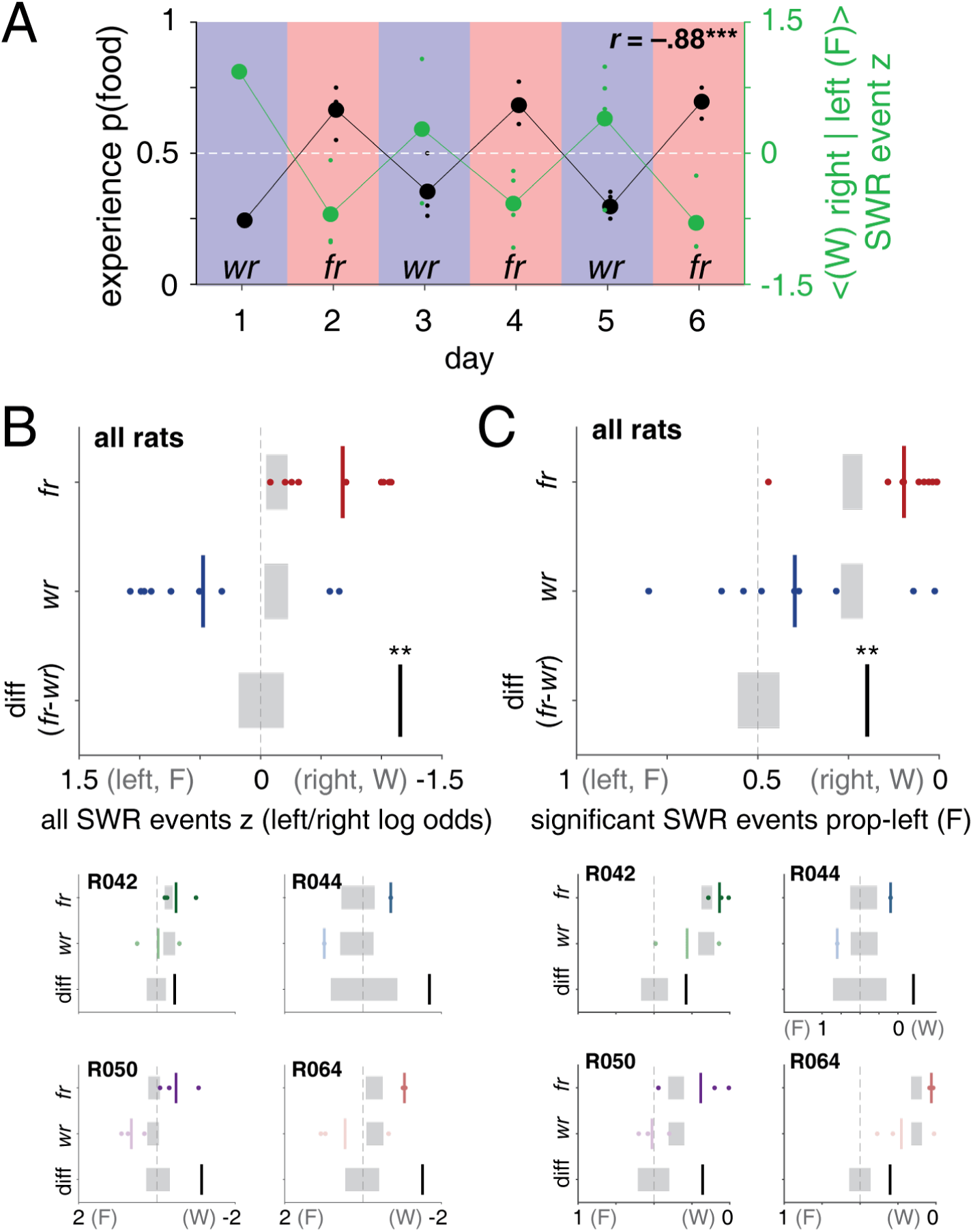
SWR content z-scored log odds based on maze arms only (i.e. excluding the central stem of the maze). Figure layout as Figure 2.

**Figure S6:**
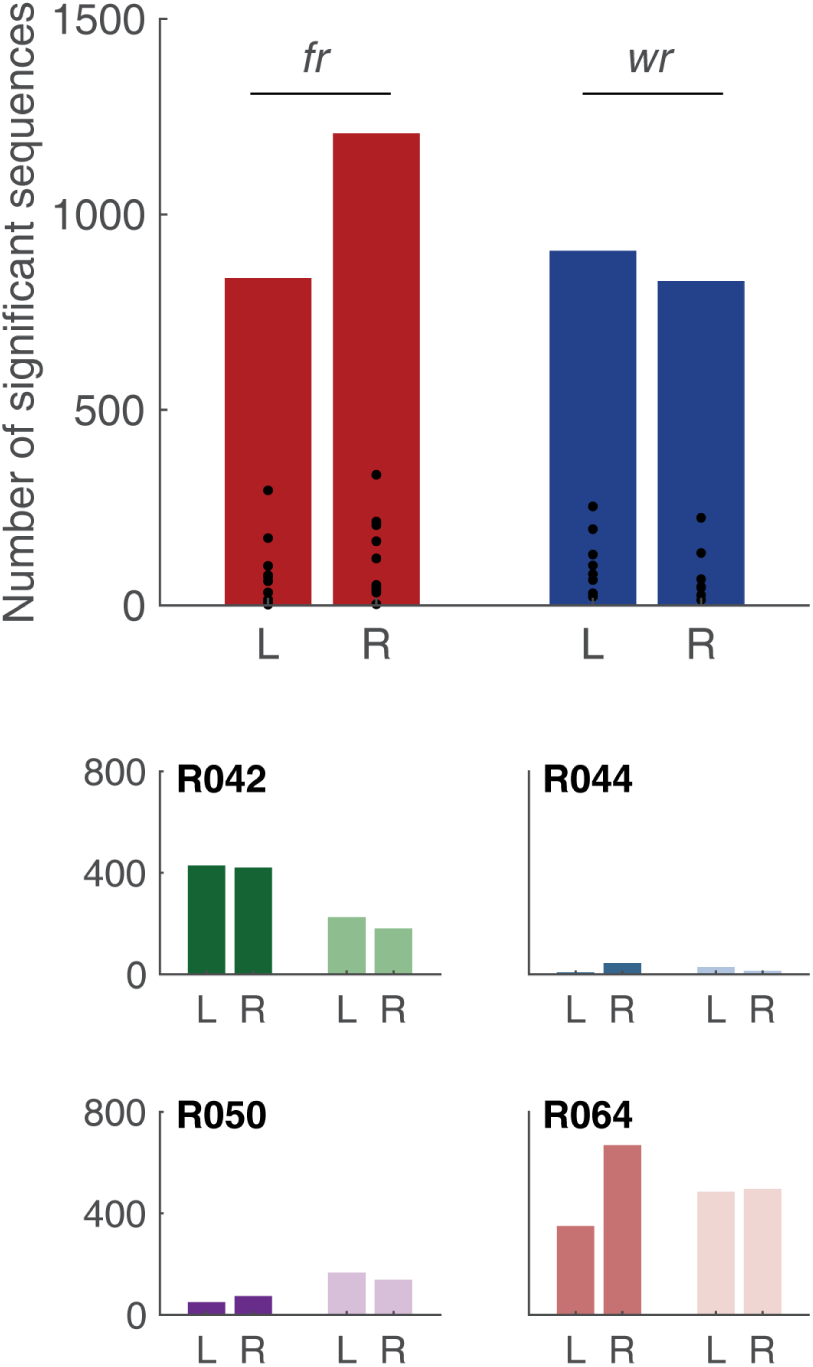
Raw SWR sequence count data for the sequence-based analysis (Figure 5).

**Figure S7:**
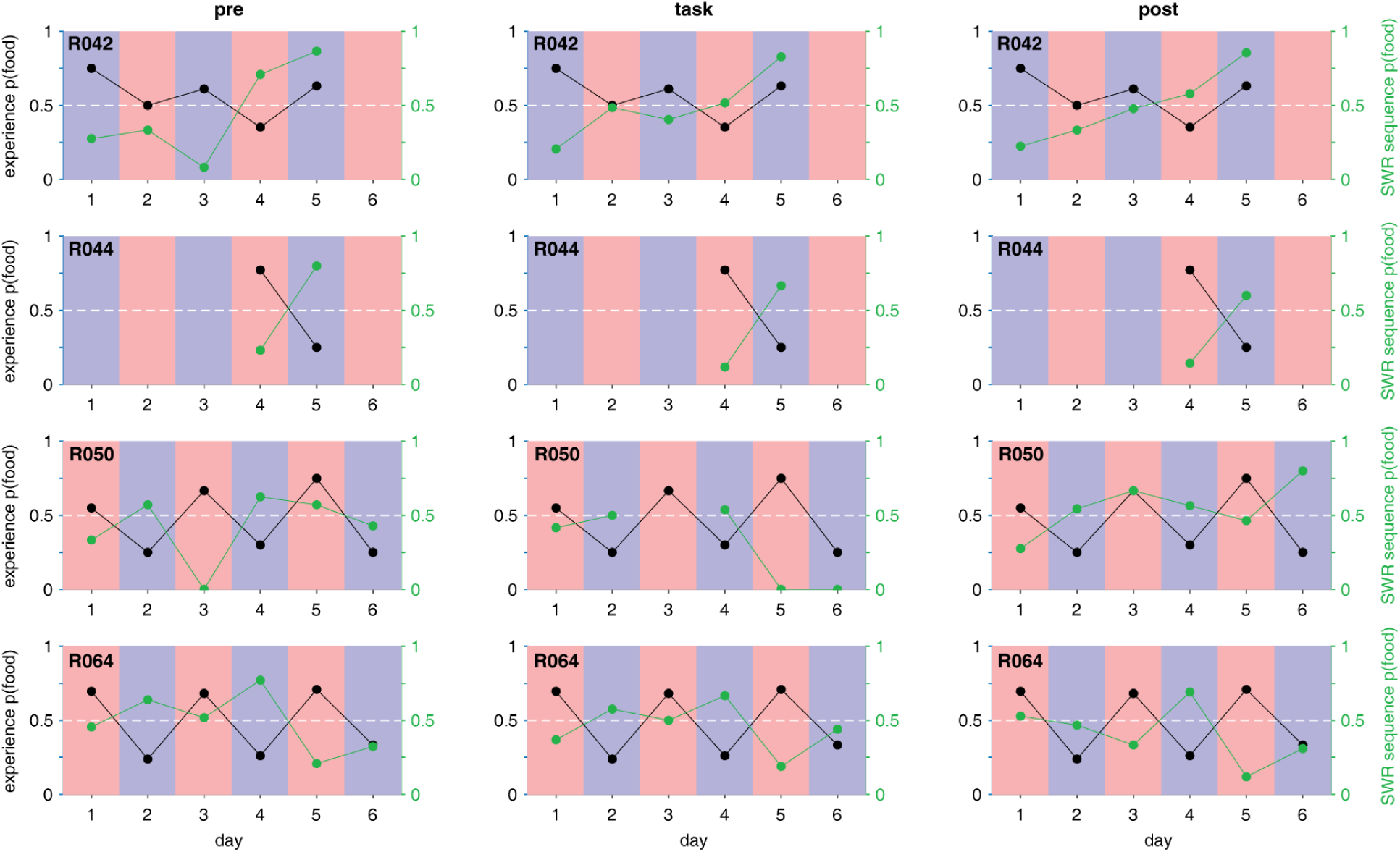
SWR sequence proportions of left arm (food) for each individual epoch and session. Layout as in Figure S4.

**Figure S8:**
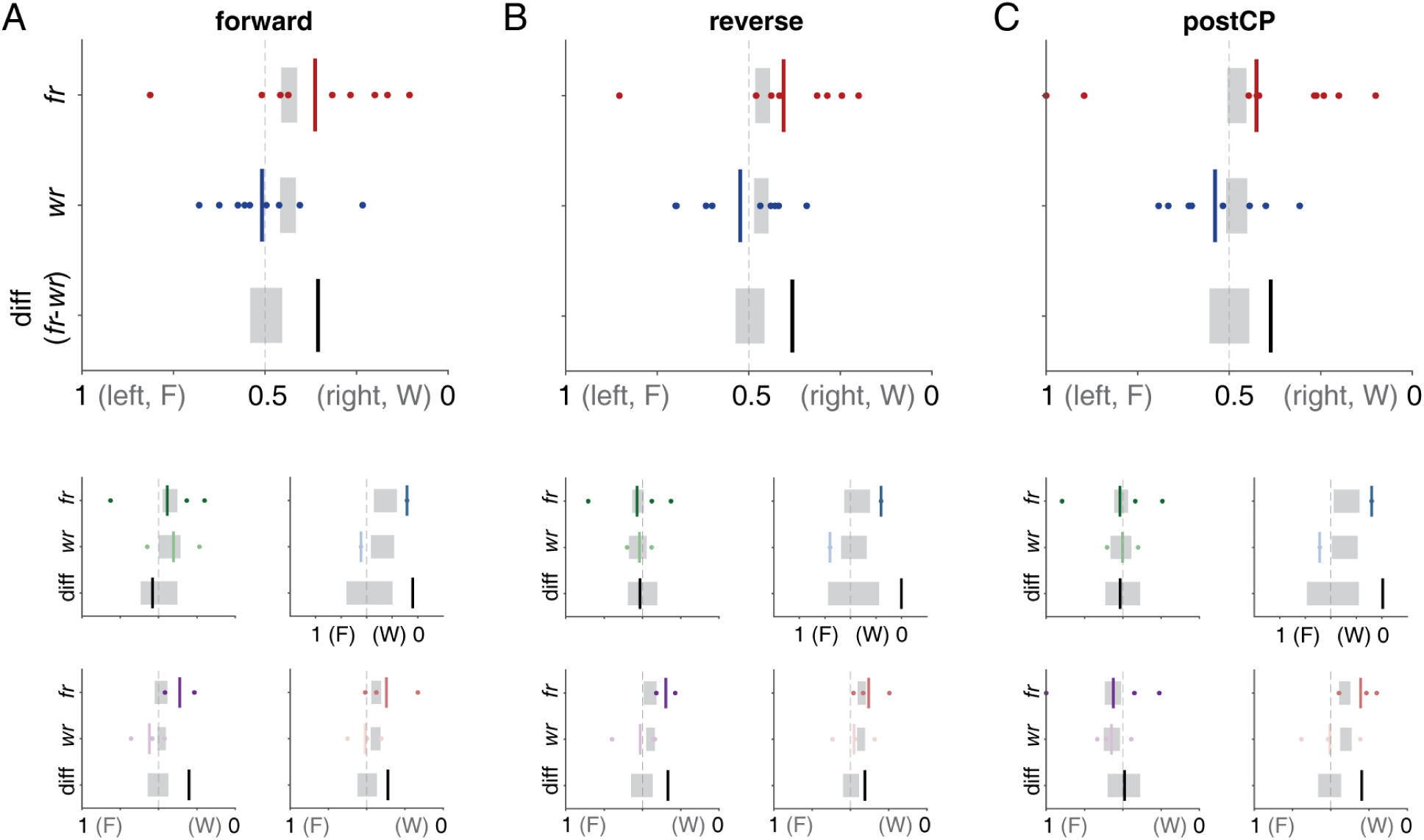
SWR sequence content (proportions left/food arm) for forward (**A**), reverse (**B**) and sequences beyond the choice point (**C**).

**Figure S9:**
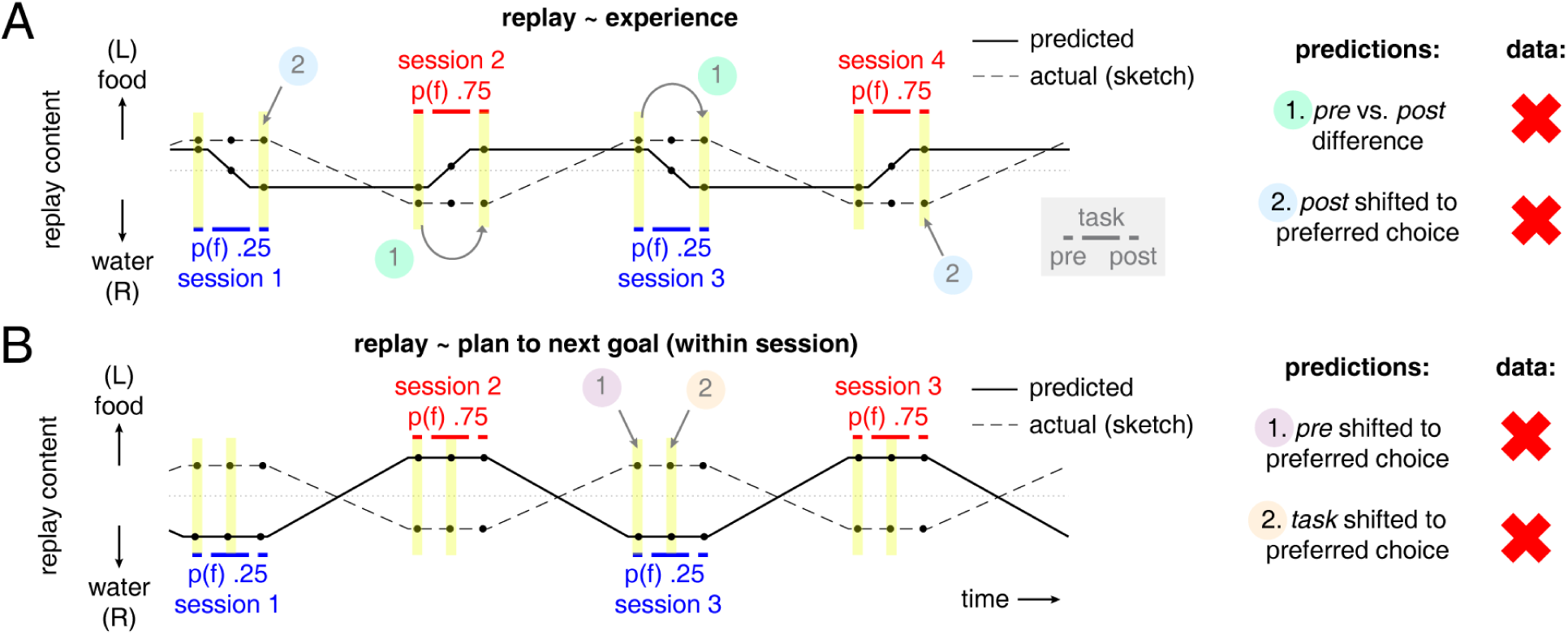
**A**: Schematic depicting the hypothesis that replay content is proportional to experience (solid line) alongside a sketch of the pattern found in the data (dashed line). The horizontal axis indicates time, including four daily recording sessions starting with a water restriction session (shown in blue on the lower left) and ending with a food restriction session (shown in red on the top right). During each recording session, experience is biased towards the restricted food type (for illustration purposes given here as a .75 probability of choosing water for water-restriction sessions, and .75 food probability for food-restriction sessions). This bias shifts predicted replay content (shown on the vertical axis) *towards* the corresponding side of the maze (right trials for water, left trials for food; note changess in the solid line (indicating predicted replay content) that occur during the task epoch of each recording session. (The size of the experience-driven changess depends on factors such as whether all experience or only within-session experience is considered, as in Figure 6c-d, but the pattern is the same.) The experience account thus predicts (1) a difference in replay content between the pre- and post-task rest periods, and (2) a bias in post-task replay content towards the recently preferred outcome. As indicated by the dashed line, neither prediction is confirmed by the data. **B**: Schematic depicting the hypothesis that replay content favors the preferred behavioral choice (or outcome). This account predicts that pre-task replay content (1) and task replay content (2) are shifted towards the preferred choice (note solid line is on the water side during water-restriction sessions, and on the food side during food-restriction sessions). As indicated by the dashed line, neither prediction is supported by the data. A variation of this account, which assumes animals can plan for the next session, correctly predicts post-task replay content, but not pre-task replay content.

**Figure S10:**
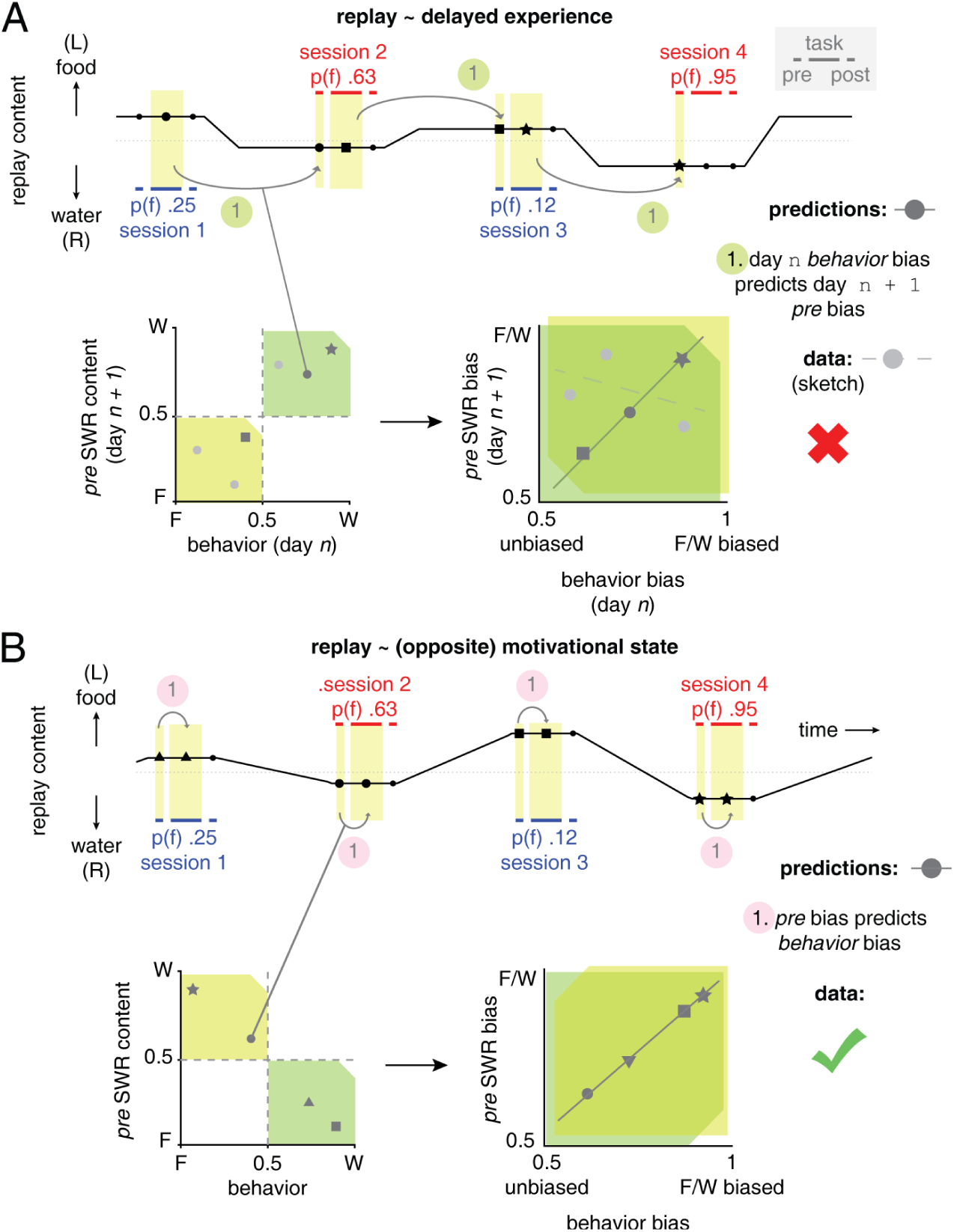
**A**: Schematic depicting the hypothesis that replay content reflects delayed experience. Although unlikely given the reported rapid effects of experience on replay content (see main text for discussion), this scenario correctly predicts no change between pre- and post-task rest, and an overall replay bias opposite the preferred outcome. However, it further predicts that the behavioral bias (preference for one side or the other, defined as *max*(*p_lef t_*, 1 *− p_lef t_*)) on day *n* predicts SWR content bias the next day: note the relatively small swing in SWR content following a relatively unbiased session (e.g. session 2 with 63% food (left) arm experience) and comparatively large swing following a strongly biased session (session 4 with 95%, top right). The two bottom panels depict the three example session data points shown (dark gray symbols), which form the predicted positive correlation; the data, depicted here schematically as light gray circles, do not exhibit such a relationship. This figure also illustrates why it is informative to compute bias scores (bottom right panel; ranging from 0.5 to 1 by using the *max* operation above) rather than using raw values (bottom left panel). This is an instance of Simpson’s Paradox, where using raw values would always show a positive correlation between day *n* behavior and *day n + 1* pre-task replay content (lower left panel): the structure of the task combined with the overall opposite bias in replay content confines the data points to the lower left and upper right quadrants. Geometrically, the bias scores align the food and water sessions to a common axis (note the reflection of the yellow and green quadrants in the lower right panel, made visible by the notch in one corner), enabling the testing of the more specific predictions shown here. **B**: If motivational state determines replay content, replay content bias during the pre-task rest period as well as other epochs should predict that session’s behavioral bias (after all, a hungrier animal would show a stronger preference for food). This prediction, illustrated in the lower two panels, is confirmed in the data. Note that again, bias scores are important in avoiding spurious results (Simpson’s Paradox). Because the comparison is within-session rather than across-session (as in **A**), the raw data are now confined to the upper left and lower right quadrants (lower left panel).

